# Seeing β-arrestin in action: The role of β-arrestins in Histamine 1 Receptor signaling

**DOI:** 10.1101/775049

**Authors:** A Pietraszewska-Bogiel, J Goedhart

## Abstract

ß-arrestins regulate G protein-coupled receptor functions by influencing their signaling activity and intracellular location. Histamine is a major chemical mediator of allergic reactions, and its action is mainly mediated by the G_q/11_- and G_i_-coupled H1R. Contrary to accumulating insights into G protein-mediated signaling downstream of H1R, very little is known about the function of ß-arrestins in H1R signaling. Here, we describe dynamic, live cell measurements of ß-arrestin recruitment upon H1R activation in HEK293TN cells. Our observations classify H1R as a class A receptor, undergoing transient interactions with ß-arrestin. To investigate the relative contributions of G proteins and ß-arrestins to H1R signaling, we use specific G protein inhibitors, as well as ß-arrestin overexpression and depletion, and quantify various signaling outcomes in a panel of dynamic, live cell biosensor assays. Overall, we link ß-arrestins to desensitization of H1R-mediated signaling and show that ERK activation downstream of endogenous (HeLa, HUVEC) or transiently expressed (HEK293TN) H1R is largely G_q_-mediated.

## INTRODUCTION

G protein-coupled or Seven transmembrane-spanning receptors (GPCRs or 7TMRs) represent the largest family of cell surface receptors. They recognize a wide variety of stimuli such as hormones, neurotransmitters, peptides, and light to regulate cellular biochemical reactions, gene expression, and cell structure and motility. Upon ligand binding, GPCRs undergo a rapid conformational change that results in their association with and activation of the heterotrimeric guanine nucleotide-binding (G) proteins, and triggering of second messenger-mediated signaling (reviewed in Hilger et. al. 2018 and Weis & Kobilka 2018). Subsequently, G protein-coupled receptor kinases (GRKs) rapidly phosphorylate specific sites within the intracellular domains of the receptor to generate high affinity arrestin binding sites (reviewed in Hilger et. al. 2018 and Komolov et al. 2018, see also Mayer et al. 2019). Out of four mammalian arrestin isoforms, two are restricted to the retina, whereas two nonvisual arrestins (arrestin2 and 3 or, respectively, ß-arrestin1 and 2) are ubiquitously expressed and bind virtually all GPCRs. Arrestin binding physically prevents GPCR-G protein interaction (although long-lasted GPCR-G protein-ßarrestin signaling complexes have also been reported, Thomsen et al. 2016), while also promoting GPCR desensitization as arrestins scaffold enzymes that degrade second messengers (Nelson et al. 2007, reviewed in Jean-Charles et al. 2017 and Gurevitch & Gurevitch 2019). Additionally, ß-arrestins function as adaptors for agonist-induced ubiquitination and endocytosis of GPCRs, regulating the fate of internalized receptors (Goodman et al. 1996, Shenoy et al. 2001, reviewed in Kang et al. 2014, Gurevich & Gurevich 2015). Receptors that interact transiently with ß-arrestins (class A receptors) undergo transient internalization, followed by recycling to the plasma membrane, whereas GPCRs that form stable complexes with ß-arrestins (class B receptors) undergo sustained internalization into endosomes (Zhang et al. 1999, Oakley et al. 2000).

Importantly, arrestins undergo conformational activation through contact with an activated GPCR (Shukla et al. 2014, Zhuo et al. 2014, Kang et al. 2015, Lee et al. 2016, Nuber et al. 2016, Eichel et al. 2018, Latorraca et al. 2018, Mayer et al. 2019), which facilitates their interactions with various effector proteins. Up to now, ß-arrestin1 (ß-arr1) and ß-arr2 were shown to scaffold multiple signaling molecules, including (but not limited to) non-receptor tyrosine kinases and mitogen-activated protein kinases (Xiao et al. 2007, reviewed in Jean-Charles et al. 2017 and Komolov et al. 2018). Therefore, arrestins also function as ligand-regulated scaffolds promoting additional (non-canonical) signaling cascades downstream from activated GPCRs (Lee et al. 2016, Nuber et al. 2016). In case of JNK3 activation, ß-arrestin stimulatory role was shown to be completely independent of G protein activation (Breitman et al. 2012)

The histamine 1 receptor (H1R) is mainly expressed in endothelium, smooth muscle cells and the central nervous system, and constitutes an important drug target in various allergic disorders, such as hay fever and urticarial and allergic rhinitis (Iriyoshi et. al. 1996, Dinh et. al. 2005, Thurmond et al. 2008). Additionally, histamine induces myometrial contractions via H1R, a process implicated in normal and preterm labor (Willets et. al. 2008 and refs. therein). H1R can activate G_q/11_ and G_i_ heterotrimeric G protein complexes (reviewed in Seifert et al. 2013, see also Adjobo-Hermans et al. 2011, van Unen et al. 2016a): G_i_ protein activation leads to inhibition of cAMP production, whereas G_q/11_ protein-mediated activation of phospholipase Cß (PLCß) results in triggering of inositol triphosphate signaling and Ca^2+^-dependent protein kinase C (PKC) pathway (Violin et al. 2003, Adjobo-Hermans et al. 2012, Ishida et al. 2014). G_q/11_ protein also plays a role in activation of a Rho GTPase, RhoA, and mitogen-activated, extracellular signal-regulated kinase (ERK) 1 and 2 upon H1R activation (Beermann et al. 2015, van Unen et al. 2015, Jain et al. 2016).

In our attempt to understand H1R signaling, we characterized recruitment of ß-arr1 and ß-arr2 upon H1R activation, as well as its implications for H1R signaling using a panel of dynamic, live cell biosensor assays. Here we show that H1R belongs to class A receptors and do not undergo pronounced internalization together with ß-arrestins. Moreover, we demonstrate redundant function of both ß-arrestin isoforms as desensitizers of G_q_-dependent signaling downstream of H1R endogenously expressed in Human Umbilical Vein Endothelial Cells (HUVEC) and HeLa cervix carcinoma cells or transiently expressed in model Human Embryonic Kidney (HEK) 293TN cells. Finally, using a specific and efficient inhibitor of G_q_ proteins, we show that H1R-mediated ERK1/2 activation in all three cell types is predominantly G_q_-dependent.

## RESULTS

### β-arrestin recruitment to the plasma membrane upon H1R activation

To begin investigating the involvement of ß-arrestins in H1R signaling, we established single cell dynamic measurements of their relocation in living cells using either a wide-field or confocal laser scanning microscope. To this end, ß-arr1 and ß-arr2 were fused to a cyan fluorescent protein, mTurquoise2 (mTQ2), at either N- or C-terminus, and H1R was fused at its C-terminus with a red fluorescent protein, mCherry (mCh), or expressed as H1R-p2A-mCherry construct (see Materials and Methods). In agreement with previous reports (e.g. Oakley et al. 2000), we observed uniform cytosolic localization of all four ß-arrestin fusions expressed in HeLa cells or HEK293TN cells (Fig. 1A-D), with ß-arr1 but not ß-arr2 showing additional nuclear localization. We did not observe any histamine-induced relocation of either ß-arrestin isoform in HeLa or HEK293TN cells when the H1R was not overexpressed. Only upon co-expression of ß-arrestin with H1R we observed histamine-induced relocation of both ß-arrestin isoforms to the cell periphery, and this process was reversed upon addition of a specific H1R antagonist, pyrilamine (PY) (Fig. 1A-D). On the wide-field microscope, this relocation was visible as a decrease of fluorescence in the cytosol and concomitant increase of fluorescence at the cell periphery (Supp. Movie 1-4). On the confocal microscope, ß-arrestin relocation was visible as a decrease of cytosolic fluorescence and concomitant recruitment of ß-arrestin to discrete puncta at the cell periphery (Fig. 1A-D, Supp. Movie 5-8). Co-localization analysis with a plasma membrane marker, Lck-mVenus, indicated that these puncta localized at or near the plasma membrane.

**Figure 1.**
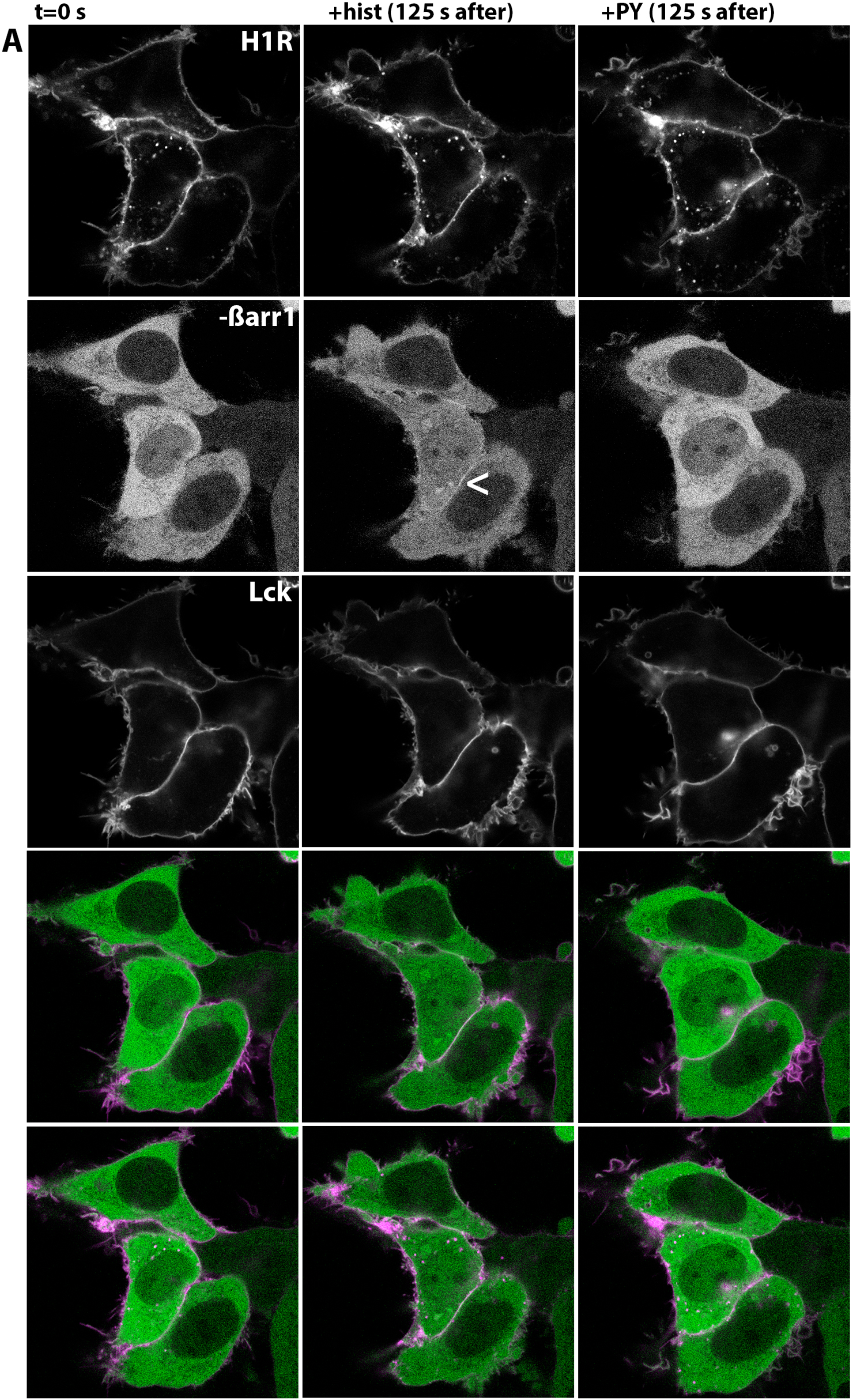
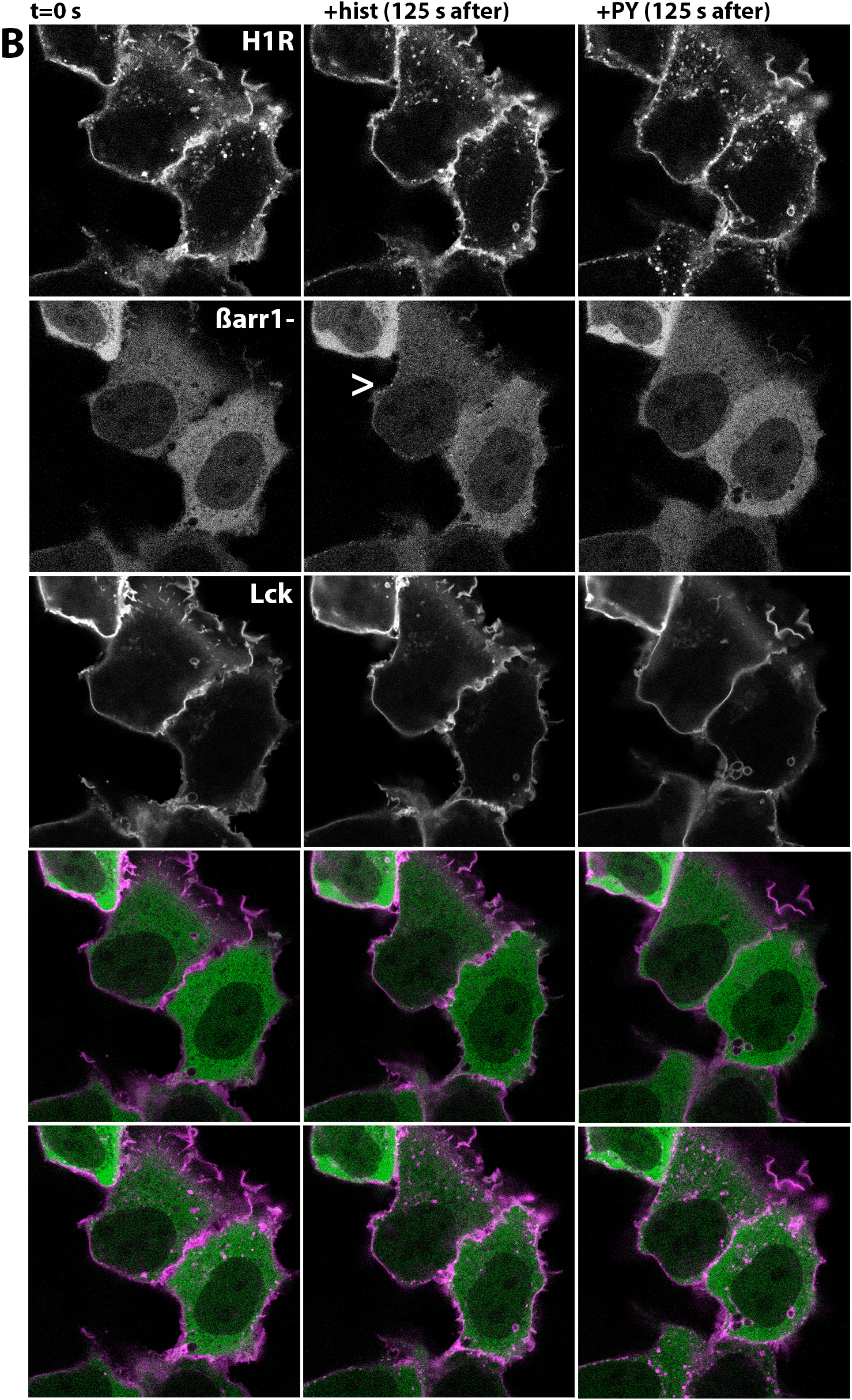
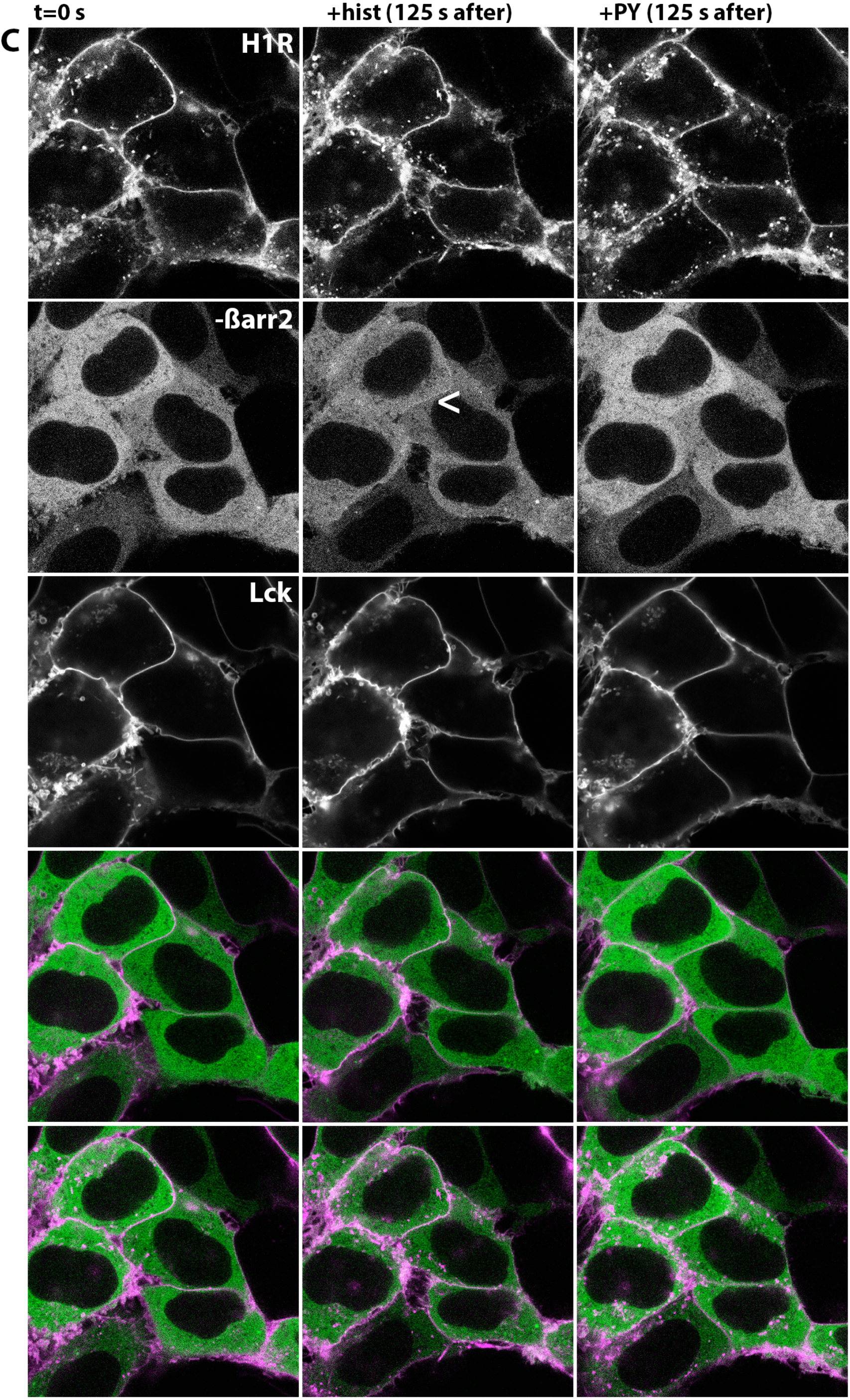
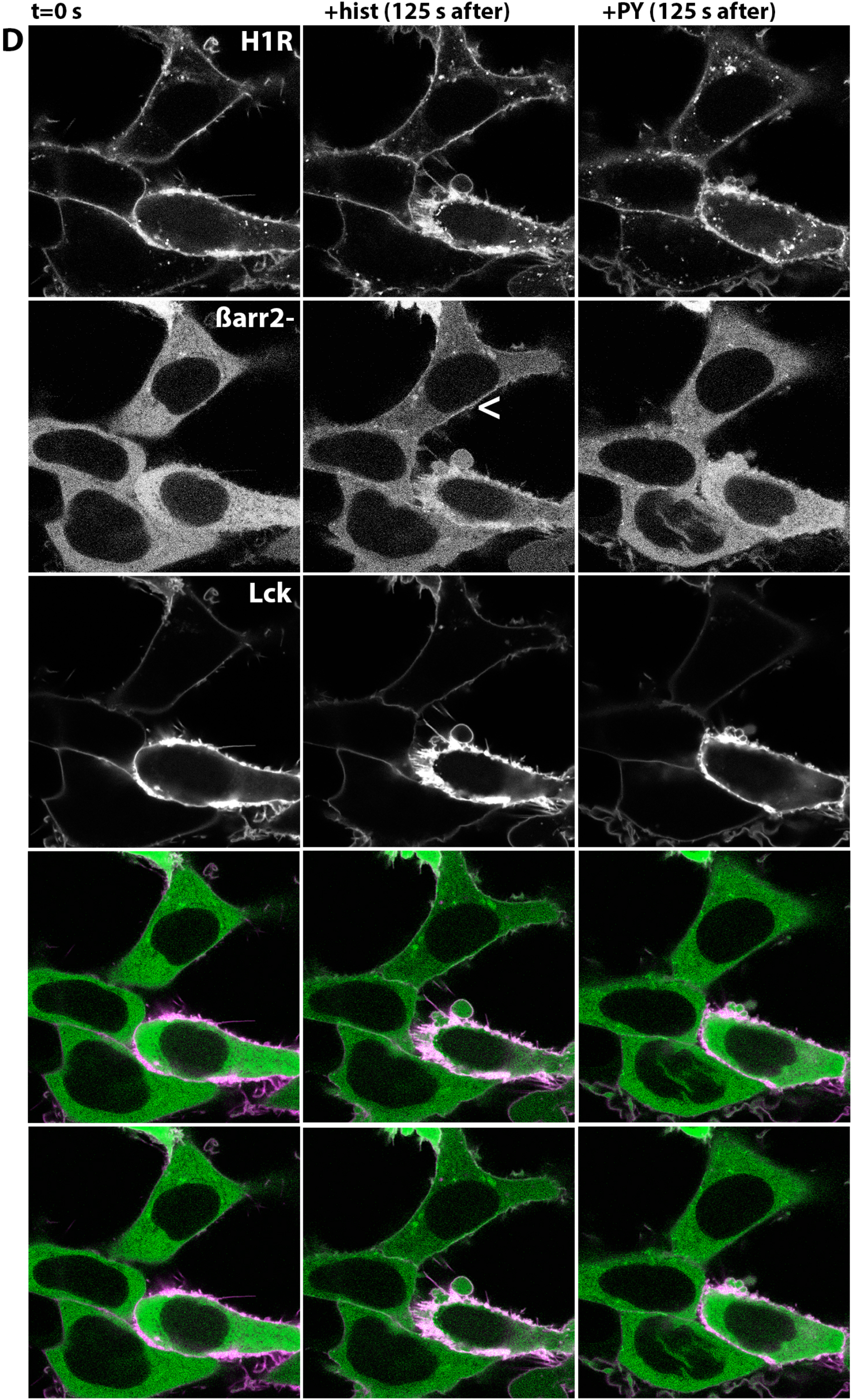
Cellular localization of ß-arrestin upon H1R activation in HEK293TN cells. A-D) Confocal images depicting subcellular localization of H1R-mCh, plasma membrane marker (Lck-mVenus) and mTQ2-ßarr1 (A), ßarr1-mTQ2 (B), mTQ2-ßarr2 (C) or ßarr2-mTQ2 (D) co-expressed in HEK293TN cells. From top to bottom: localization of H1R-mCh; localization of ßarr; localization of Lck-mVenus; merged image of Lck (set to magenta) and ßarr (set to green) localization, where the shared localization is depicted in white; merged image of H1R (set to magenta) and ßarr (set to green) localization, where the shared localization is depicted in white. From left to right: localization prior histamine stimulation, 125 s after addition of 100 μM histamine, and 125 s after addition of 10 μM pyrilamine. Arrowheads point to the histamine-induced localization of ß-arrestin. The size of the images is 60 × 60 μm.

GPCRs can be functionally divided into two classes based on the character and outcomes of their interaction with ß-arrestins (Zhang et al. 1999, Oakley et al. 2000). In order to qualitatively classify H1R to either class A or class B, we compared H1R-mediated, beta2-adrenergic receptor (ß2AR, class A)-mediated, and angiotensin II type 1 receptor (AT1_A_R, class B)-mediated relocation of ß-arr2. To this end, AT1_A_R and ß2AR were fused at their C-terminus with mCherry, and co-expressed together with Lck-mVenus and ßarr2-mTQ2 in HEK293TN cells. We observed robust agonist-induced relocation of ß-arr2 in cells co-expressing the receptor (Fig. 2A&B, Supp. Movie 9&10) but not in cells expressing only Lck-mVenus and ßarr2-mTQ2 (data not shown). ß2AR activation with isoproterenol resulted in a decrease of cytosolic fluorescence and concomitant recruitment of ß-arrestin to discrete puncta at the cell periphery. Co-localization analysis with Lck-mVenus indicated plasma membrane localization of these puncta. On the contrary, AT1_A_R activation with angiotensin II (AngII) resulted in robust relocation of ßarr2, first to the plasma membrane (as indicated by its co-localization with Lck-mVenus), and subsequently to stable intracellular puncta, most likely endosomal compartments. Importantly, we observed extensive co-localization of AT1_A_R with ßarr2 at this intracellular compartment.

**Figure 2.**
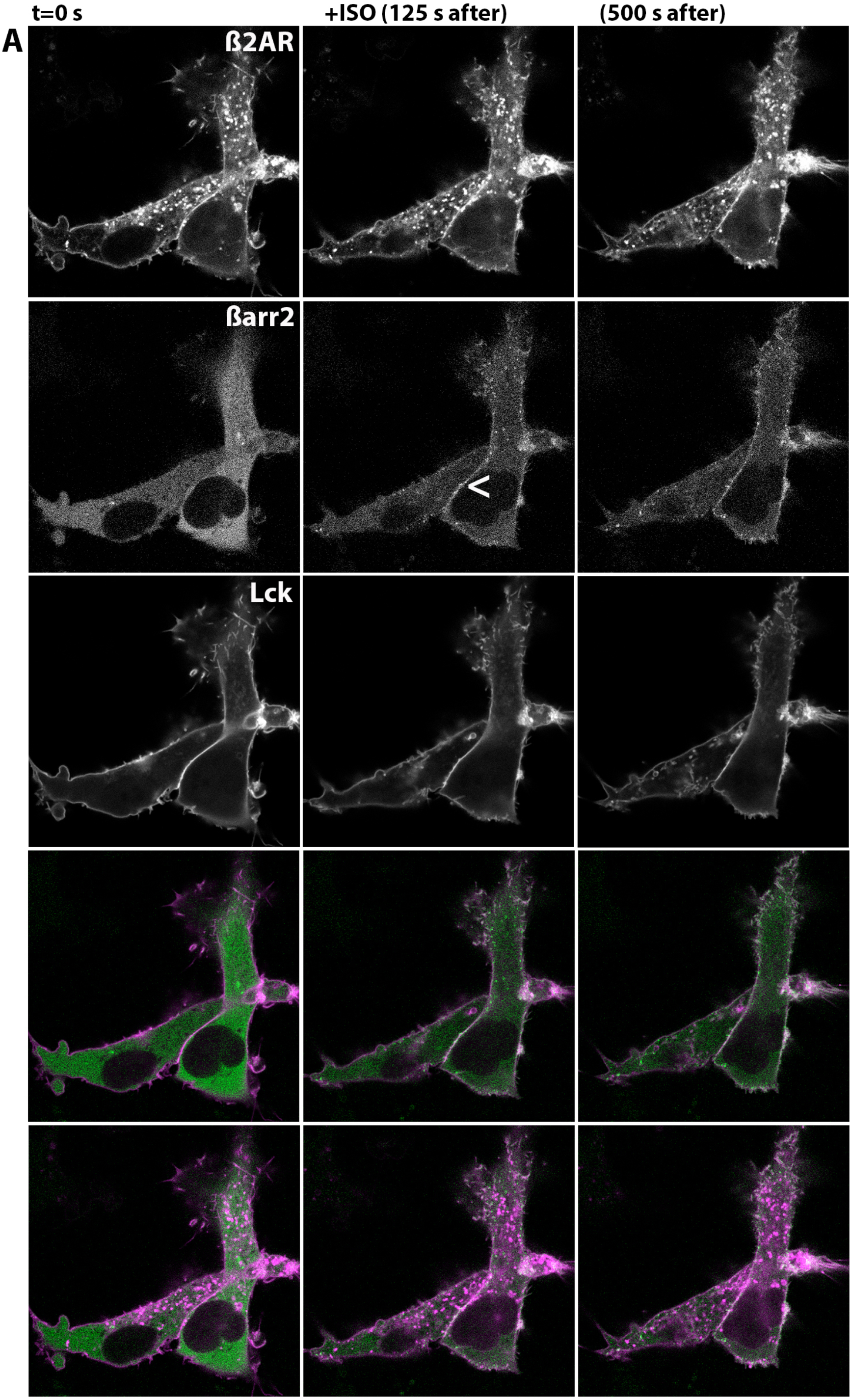
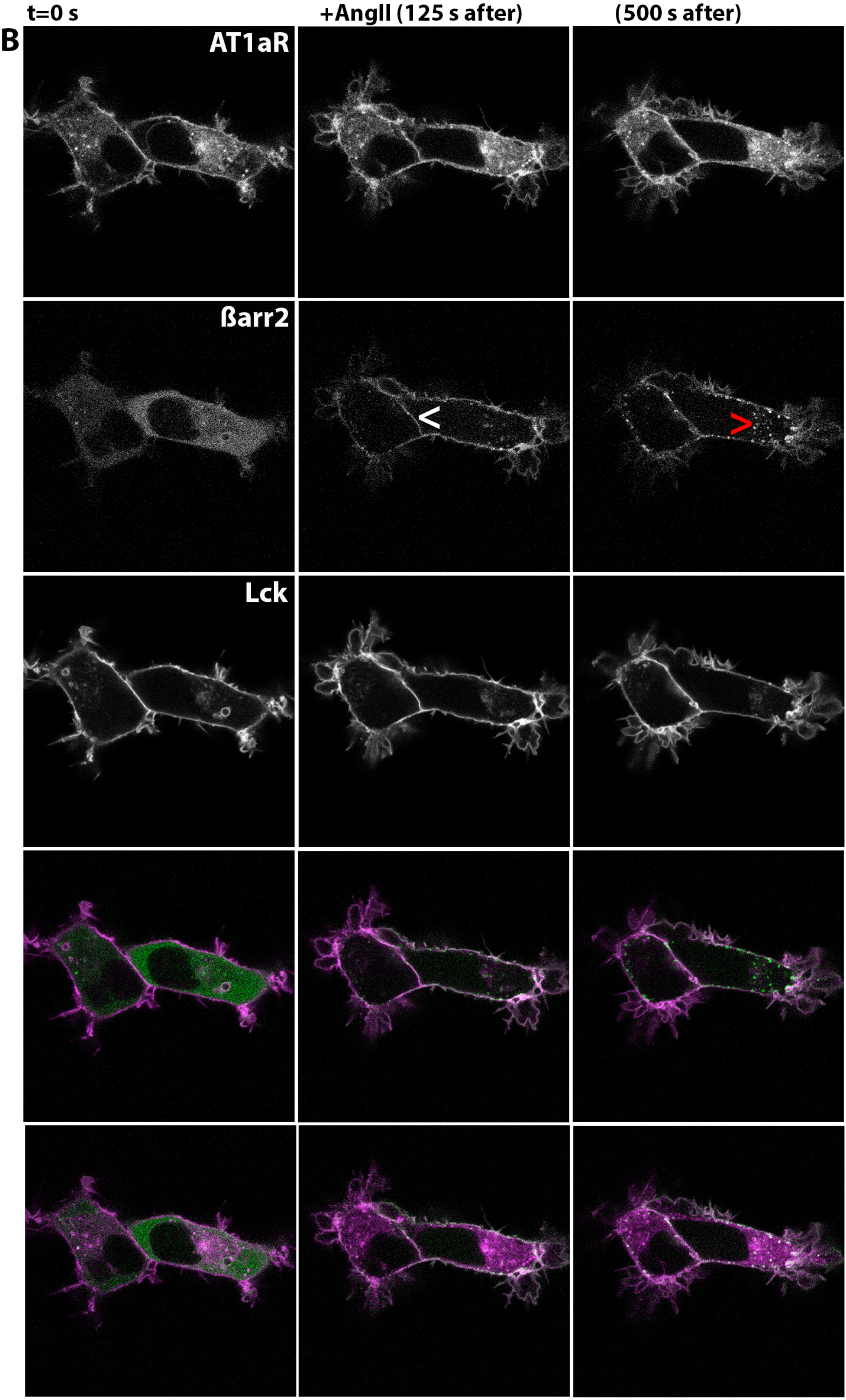
Cellular localization of ß-arrestin upon ß2AR and AT1_A_R activation in HEK293TN cells. A) Confocal images depicting subcellular localization of ß2AR-mCh, Lck-mVenus and ßarr2-mTQ2 co-expressed in HEK293TN cells. From top to bottom: localization of ß2AR-mCh; localization of ßarr2-mTQ2; localization of Lck-mVenus; merged image of Lck (set to magenta) and ßarr2 (set to green) localization, where the shared localization is depicted in white; merged image of ß2AR (set to magenta) and ßarr2 (set to green) localization, where the shared localization is depicted in white. From left to right: localization prior, 125 s after, and 500 s after 10 μM isoproterenol stimulation. Arrowhead points to the ISO-induced localization of ß-arrestin at or near the plasma membrane. The size of the images is 60 × 60 μm. B) Confocal images depicting subcellular localization of AT1_A_R-mCh co-expressed together with Lck-mVenus and ßarr2-mTQ2 in HEK293TN cells. From top to bottom: localization of AT1_A_R-mCh; localization of ßarr2-mTQ2; localization of Lck-mVenus; merged image of Lck (set to magenta) and ßarr2 (set to green) localization, where the shared localization is depicted in white; merged image of AT1_A_R (set to magenta) and ßarr2 (set to green) localization, where the shared localization is depicted in white. From left to right: subcellular localization prior, 125 s after, and 500 s after AngII II stimulation. Arrowheads point to the AngII-induced localization of ß-arrestin at or near the plasma membrane (white) or in the intracellular compartment (red). The size of the images is 60 × 60 μm.

To quantify ß-arrestin recruitment upon H1R activation, we co-expressed H1R-mCh together with ßarr1-mTQ2 or ßarr2-mTQ2 in HEK293TN cells, recorded ß-arrestin relocation on the confocal microscope, and then measured histamine-induced decrease of cyan fluorescence in the cytosol. We measured similar kinetics of relocation for both ß-arrestin isoforms: half-time (t_1/2_) of 50 s for mTQ2-ßarr2 and ßarr2-mTQ2 relocation, and 75 s for mTQ2-ßarr1 and ßarr1-mTQ2 (Fig. 3A&B). However, relocation of ß-arr2 fusions was more robust (32% drop of intensity) in comparison to relocation of ß-arr1 fusions (20% drop of intensity). We did not measure any change in the abundance of nuclear pool of ß-arr1 (Fig. 3A). Interestingly, for both ß-arrestin isoforms, the increase of cytosolic fluorescence upon pyrilamine addition exceeded the baseline levels measured prior histamine stimulation.

**Figure. 3.**
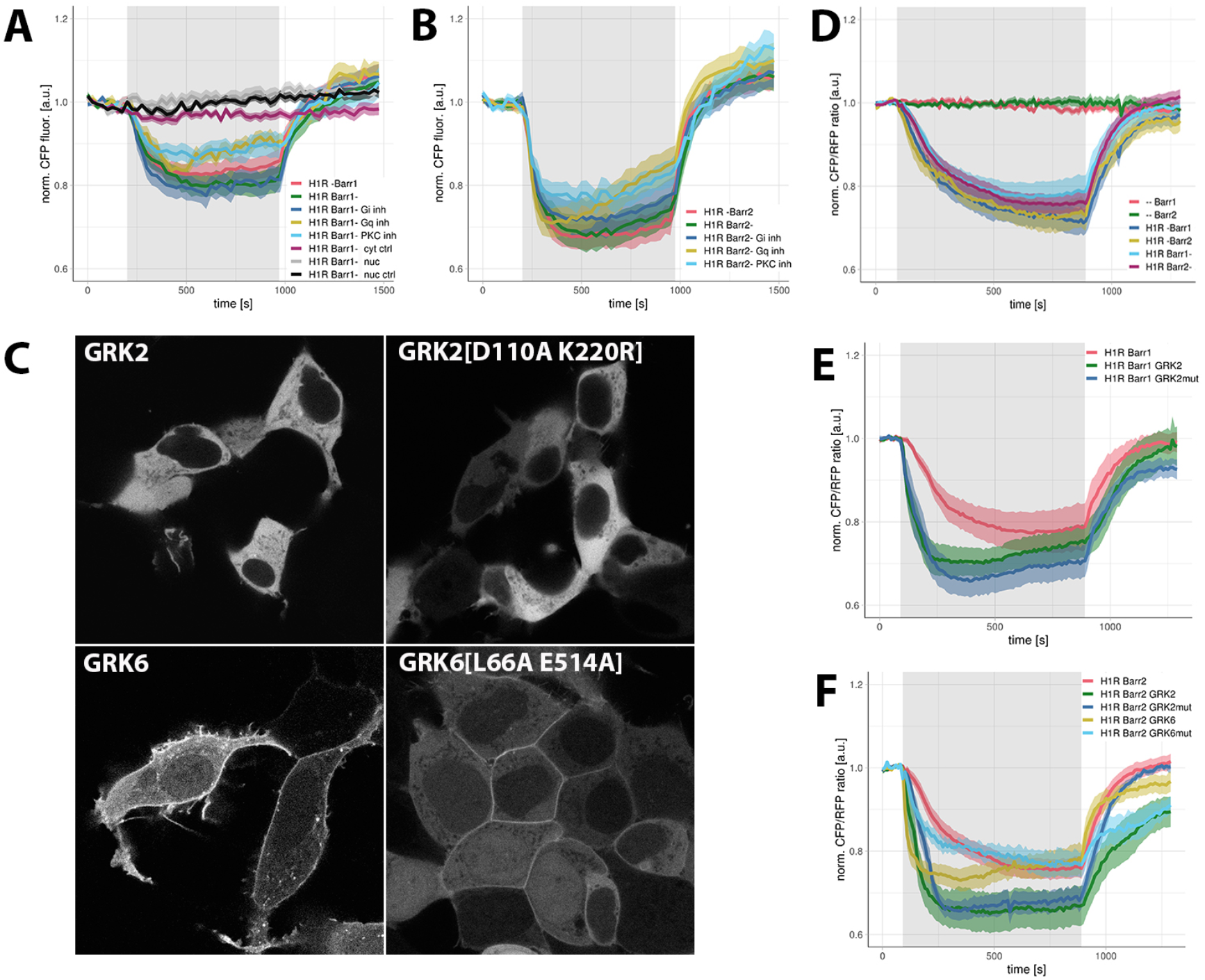
Quantification of ß-arrestin recruitment upon H1R activation. A-B) Confocal measurements of ß-arr1 (A) or ß-arr2 (B) relocation. Time traces show the average CFP fluorescence (±95% confidence intervals) measured in the cytosol of HEK293TN cells, except for the grey and black traces which show the average CFP fluorescence (±95% confidence intervals) measured in the nucleus. Vehicle (ctrl) or 100 μM histamine was added at 90 s and 10 μM PY at 890 s; grey boxes mark the duration of histamine stimulation. A) HEK293TN cells co-expressing: H1R-mCh and mTQ2-ßarr1 (-ßarr1); H1R-mCh and ßarr1-mTQ2 (ßarr1-) and untreated, treated with Gα_i_ inhibitor (PTX), Gα_q_ inhibitor (FR900359) or PKC inhibitor (Ro31-8425). B) HEK293TN cells co-expressing: H1R-mCh and mTQ2-ßarr2 (-ßarr2); H1R-mCh and ßarr2-mTQ2 (ßarr2-) and untreated, treated with PTX, FR900359 or Ro31-8425. C) Confocal images depicting subcellular localization of GRK2-sYFP2, GRK2[D110A K220R]-sYFP2, GRK6-sYFP2, and GRK6[L66A E514A]-sYFP2 expressed in HEK293TN cells. The size of the images is 105 × 105 μm. D-F) Wide-field measurements of ß-arrestin relocation. Time traces show the average ratio change of CFP/RFP fluorescence (±95% confidence intervals) measured in the cytosol of HeLa cells. 100 μM histamine or vehicle (--) was added at 90 s and 10 μM PY at 890 s; grey boxes mark the duration of histamine stimulation. HeLa cells: D) expressing ßarr1-mTQ2 or ßarr2-mTQ2 alone or co-expressing H1R-p2A-mCh together with ßarr1-mTQ2, mTQ2-ßarr1, ßarr2-mTQ2 or mTQ2-ßarr2; E) co-expressing H1R-p2A-mCh and ßarr1-mTQ2 only or together with GRK2-sYFP2 or GRK2[D110A K220R]-sYFP2; F) co-expressing H1R-p2A-mCh and ßarr2-mTQ2 only or together with GRK2-sYFP2, GRK2[D110A K220R]-sYFP2, GRK6-sYFP2 or GRK6[L66A E514A]-sYFP2.

Therefore, H1R activation resulted in relocation of both ß-arrestin isoforms to the plasma membrane, and this ß-arrestin recruitment resembled more the recruitment by a class A receptor rather than by a class B GPCR.

### Requirements for β-arrestin recruitment upon H1R activation

Subsequently, we investigated whether G protein activation is required for ß-arrestin recruitment upon H1R activation. To this end, we measured kinetics of ß-arrestin relocation in HEK293TN cells in the absence and presence of a specific Gα_q_ inhibitor (FR900359, Schrage et al. 2015) or a specific Gα_i_ inhibitor, pertussis toxin (PTX). We measured similar kinetics of relocation for ßarr1-mTQ2 and ßarr2-mTQ2 in the absence or presence of G_i_ protein activation, whereas ß-arrestin relocation in the absence of G_q_ protein activation was more transient, with part of the ß-arrestin fusion proteins returning to the cytosol even before the addition of pyrilamine (Fig. 3A&B). As H1R was shown to undergo PKC-dependent phosphorylation on Ser396 and Ser398 (Fujimoto et al. 1999, Horio et al. 2004), we also investigated the role of PKC in ß-arrestin recruitment using a specific PKC inhibitor, Ro31-8425. Relocation of both ß-arrestin isoforms was decreased in the absence of PKC activation (Fig. 3A&B): we measured 13% and 25% drop of cytosolic intensity, respectively, for ßarr1-mTQ2 and ßarr2-mTQ2.

In addition to PKC-mediated phosphorylation, H1R was shown to be phosphorylated by GRK2, and to a lesser extent by GRK5 and GRK6, in uterine myometrial cells and HEK293 cells (Iwata et al. 2005, Willets et al. 2008). Unfortunately, phosphorylation sites undergoing GRK-dependent phosphorylation in H1R are currently unknown, which occludes the generation of a phosphorylation-deficient H1R mutant. Therefore, we investigated GRK role by measuring the effect of GRK2 and GRK6 overexpression on histamine-induced ß-arrestin relocation in HeLa cells. As a reference, we used two GRK mutants postulated to be catalytically impaired: GRK2[D110A K220R] (Iwata et al. 2005) and GRK6[L66A E514A]; the latter mutant corresponds to GRK5[L66A E514A] mutant described by Baameur et al. (2010) (see also Discussion). To visualize GRK expression, WT and mutated GRK2 and GRK6 were fused at their C-terminus with a yellow fluorescent protein, sYFP2, and their subcellular localization was analyzed in HEK293TN cells. Both GRK2-sYFP2 and GRK2[D110A K220R]-sYFP2 fusions were localized exclusively in the cytosol, whereas GRK6-sYFP2 and GRK6[L66A E514A]-sYFP2 fusions were localized predominantly at the plasma membrane, with additional localization in the cytosol and nucleus (Fig. 3C).

Then, we co-expressed H1R-p2A-mCh together with ßarr1-mTQ2 or ßarr2-mTQ2 in HeLa cells, and measured ß-arresting relocation in the absence and presence of GRK overexpression. In the former situation, we measured t_1/2_ of 140 s and 25% drop in cytosolic intensity for both ßarr1-mTQ2 and ßarr2-mTQ2 (Fig. 3D). However, as those measurements were done on a wide-field microscope, the amplitude of ß-arrestin relocation upon H1R activation in HeLa and HEK293TN cells cannot be directly compared. In HeLa cells co-expressing H1R-p2A-mCh, GRK2-sYFP2 and ßarr1 or ßarr2 fusions, we measured t_1/2_ of 35 s and 30% drop of intensity with regard to ß-arr1 relocation, and 55 s and 35% drop of intensity with regard to ß-arr2 relocation (Fig. 3E&F). Surprisingly, relocation of both ß-arrestin isoforms was similarly enhanced in cells co-expressing H1R-p2A-mCh and GRK2[D110A K220R]-sYFP2: t_1/2_ of 50 s and 90 s measured for ßarr1-mTQ2 and ßarr2-mTQ2, respectively (Fig. 3E and F). The effect of GRK6 overexpression was only tested with regard to ß-arr2 relocation: we measured t_1/2_ of 15 s for ßarr2-mTQ2 in cells overexpressing GRK6-sYFP2, and 80 s in cells overexpressing GRK6[L66A E514A]-sYFP2 (Fig. 3F).

Taken together, we showed that ß-arrestin relocation upon activation of H1R did not rely on G_q_ or G_i_ activation. Instead, our data on the GRK role indicated, although indirectly, a likely dependence of ß-arrestin recruitment on the H1R phosphorylation.

### β-arrestin-mediated desensitization of H1R signaling via Gq protein

We then characterized the role of ß-arrestin in desensitization of H1R-mediated signaling, looking at the dynamics of the response in the continuous presence of the agonist. Specifically, we measured G_q_-mediated, PLCß-dependent degradation of PtdIns(4,5), PIP2, in the plasma membrane, which can be observed in living cells as relocation of the PIP2 sensor (mTQ2-PHM, Adjobo-Hermans et al. 2012) from the plasma membrane to the cytosol. First, we studied the effect of ß-arrestin overexpression on histamine-induced PIP2 degradation. To this end, we fused ß-arr1 and ß-arr2 at their C-terminus with sYFP2 (ß-arrestin fusions to sYFP2 showed the same subcellular localization as the respective fusions to mTQ2, data not shown) Then, we co-expressed mTQ2-PHM and H1R-p2A-mCh alone or together with either ßarr1-sYFP2 or ßarr2-sYFP2 in HeLa cells, and measured PIP2 sensor relocation. We observed a pronounced and persistent increase in cytosolic localization of mTQ2-PHM upon histamine stimulation that was reversed with pyrilamine (Fig. 4A). Co-expression of either ßarr1-sYFP2 or ßarr2-sYFP2 resulted in a faster replenishment of the PIP2 sensor at the plasma membrane, although the cytosolic levels of mTQ2-PHM returned to baseline level still only upon addition of PY.

**Figure. 4.**
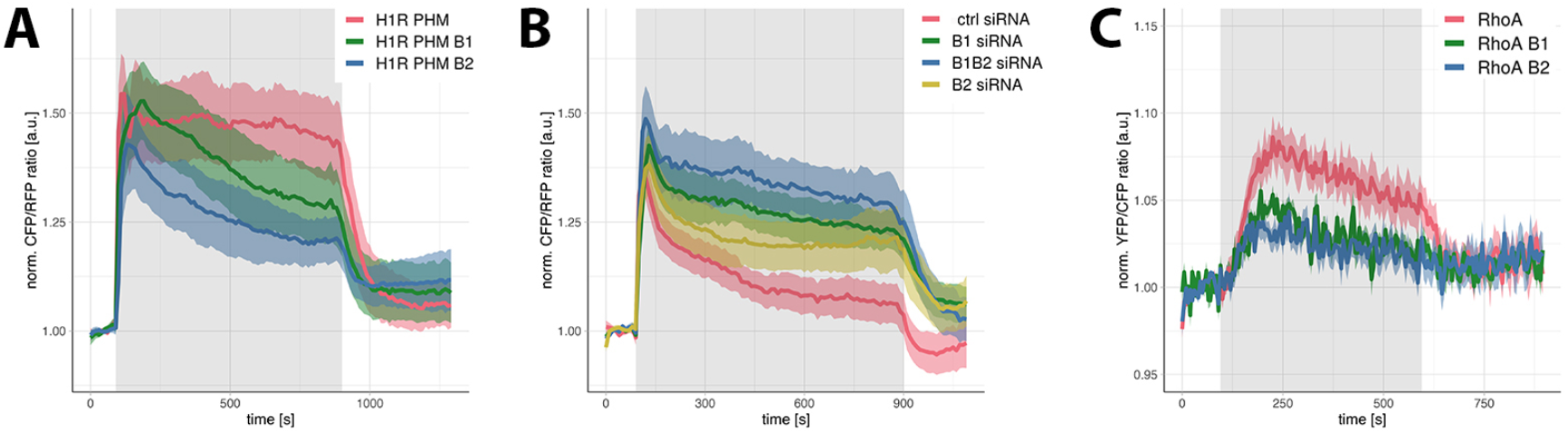
Role of ß-arrestins in G_q_-mediated signaling downstream of H1R. A) The effect of ß-arrestin overexpression on histamine-induced PLCß activation in HeLa cells. Time traces show the average CFP/RFP ratio change (±95% confidence intervals) in cells expressing H1R-p2A-mCh and PIP2 reporter (mTQ2-PHM) or cells expressing H1R-p2A-mCh, mTQ2-PHM, and either ßarr1-sYFP2 or ßarr2-sYFP2. 100 μM histamine was added at 90 s and 10 μM pyrilamine at 890 s; the grey box marks the duration of histamine stimulation. B) The effect of ß-arrestin knockdown on histamine-induced PLC activation in HEK293TN cells. Time traces show the average CFP/RFP ratio change (±95% confidence intervals) in cells expressing H1R-p2A-mCh and mTQ2-PHM, that were first transfected with ctrl siRNA, ß-arr1 siRNA, ß-arr2 siRNA or simultaneously ß-arr1 and ß-arr2 siRNA constructs. 100 μM histamine was added at 90 s and 10 μM PY at 890 s; the grey box marks the duration of histamine stimulation. C) The effect of ß-arrestin overexpression on histamine-induced RhoA activation in HUVEC cells. Time traces show the average YFP/CFP ratio change (±95% confidence intervals) in cells expressing DORA-RhoA reporter (RhoA) alone or expressing DORA-RhoA and either ßarr1-mCh or ßarr2-mCh. 100 μM histamine was added at 95 s and 10 μM PY at 595 s; the grey box marks the duration of histamine stimulation.

Subsequently, we studied the effect of ß-arrestin knockdown on histamine-induced PIP2 degradation in HEK293TN cells. To this end, we transfected cells with ctrl siRNA, ß-arr1 siRNA, ß-arr2 siRNA or simultaneously ß-arr1 and ß-arr2 siRNA constructs (for the efficiency of ß-arr1 and/or ß-arr2 knockdown, see Supp. Fig. 3). Twenty-four hours later, the cells were transfected with H1R-p2A-mCh and mTQ2-PHM constructs, and relocation of the PIP2 sensor was measured after additional 24 h. Histamine stimulation of cells transfected with ctrl siRNA resulted in a pronounced increase in cytosolic localization of mTQ2-PHM, although it showed a faster decay in comparison to the response of HeLa cells. Silencing of either ß-arr1 or ß-arr2 resulted in more persistent cytosolic localization of mTQ2-PHM, with simultaneous silencing of both ß-arrestin isoforms showing an additive effect (Fig. 4B).

Additionally, we studied ß-arrestins’ role in signaling mediated by endogenous H1R in HUVEC cells. We chose to measure the effect of ß-arrestin overexpression on RhoA activation using DORA-RhoA FRET sensor (Reinhard et al. 2016). To this end, we fused ß-arrestins at their C-terminus with mCherry (see Supp. Fig. 4 for their subcellular localization), and co-expressed the resulting ßarr1-mCh or ßarr2-mCh constructs together with DORA-RhoA sensor. We measured reproducible histamine-induced DORA-RhoA signals that were reversed upon pyrilamine addition (Fig. 4C). Co-expression of either ß-arr1 or ß-arr2 diminished this FRET signal to a similar extent.

Taken together, we demonstrated a largely redundant role of ß-arr1 and ß-arr2 in desensitization of G_q_-mediated signaling downstream of H1R.

### Histamine-mediated activation of ERK1/2

Finally, we characterized ERK1/2 activation downstream of endogenous H1R expressed in HeLa and HUVEC cells. We measured ERK1/2 activation separately in the cytosol and in the nucleus by employing, respectively, EKARcyt and EKARnuc sensors (Harvey et al. 2008). We monitored EKAR signals for more than 30 minutes to measure both fast and late phases of ERK-mediated phosphorylation.

Stimulation of HeLa cells with histamine activated the cytosolic and nuclear pool of ERK1/2 with similar kinetics (Fig. 5A&B): both EKAR signals increased gradually to achieve maxima approximately 380 s after stimulation, and then slowly decayed. Additionally, we observed a transient drop of EKAR signal (very small for EKARnuc, 30% of maximal value for EKARcyt signal) immediately after histamine addition. Inhibition of Gα_q_ with FR900359 completely abolished both EKAR signals, whereas inhibition of Gα_i_ with pertussis toxin partially diminished EKARnuc but not EKARcyt signal. Finally, as G_q_ protein-dependent ERK1/2 activation can be mediated by PKC (reviewed in DeWire et al. 2007), we measured EKAR signals in cells incubated with Ro31-8425. Interestingly, PKC inhibitor strongly diminished but did not completely inhibit either EKAR signal (Fig. 5A&B).

**Figure 5.**
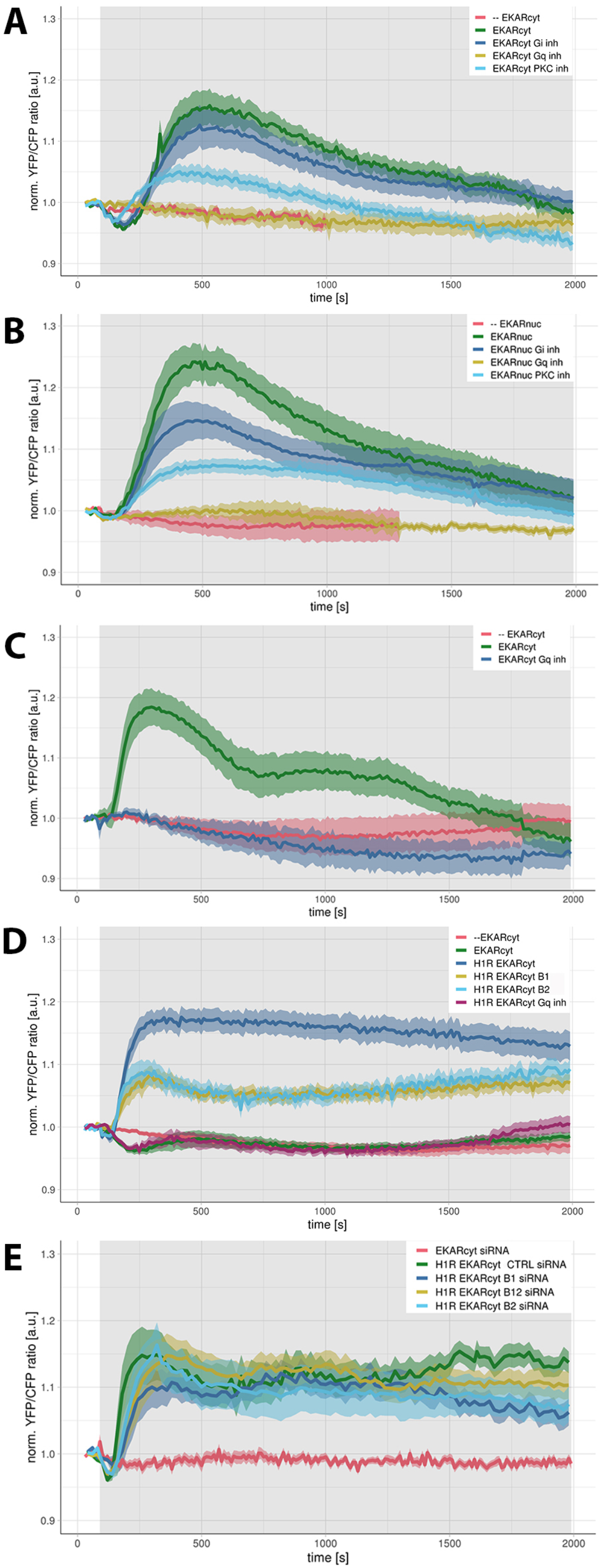
H1R-mediated activation of ERK1/2. A-E) Time traces show the average ratio change of YFP/CFP fluorescence (±95% confidence intervals). Vehicle (--) or 100 μM histamine was added at 90 s; grey boxes mark the duration of histamine stimulation. A-B) HeLa cells expressing EKARcyt (A) or EKARnuc (B) sensor were untreated, treated with Gα_i_ inhibitor (PTX), Gα_q_ inhibitor (FR900359) or PKC inhibitor (Ro31-8425), C) HUVEC cells expressing EKARcyt sensor were untreated or treated with FR900359, D) HEK293TN cells: co-expressing EKARcyt together with H1R-p2A-mCh were untreated or treated with FR900359; co-expressing EKARcyt, H1R, and ßarr1-mCh or ßarr2-mCh, E) HEK293TN cells co-expressing EKARcyt sensor together with H1R-p2A-mCh, that were first transfected with ctrl siRNA, ß-arr1 siRNA, ß-arr2 siRNA or simultaneously ß-arr1 and ß-arr2 siRNA constructs. Red trace shows the response of cells expressing only the EKARcyt reporter, that were first transfected with siRNA constructs (data from ctrl, ßarr1, and ßarr2 cells were pulled together).

Histamine stimulation of HUVEC cells also resulted in robust activation of the cytosolic pool of ERK1/2 (Fig. 5C, activation of nuclear pool of ERK1/2 not tested). In comparison with EKARcyt signals measured in HeLa cells, ERK1/2 activation in HUVEC cells showed slightly faster kinetics but lower amplitude, and decayed in biphasic mode: upon reaching the maximum at 200 s after histamine addition, the signal first dropped by 50% within next 350 s, and then stayed constant for approximately 500 s before gradually dropping to baseline approximately 30 min after histamine addition. Similar to HeLa cells, inhibition of Gα_q_ with FR900359 completely abolished histamine-induced EKARcyt signal in HUVEC cells (Fig. 5C). Furthermore, we wanted to investigate the role of ß-arrestins in ERK1/2 activation in HUVEC cells using both overexpression and knockdown strategies. However, expression of multiple constructs at sufficient levels turned out to be notoriously difficult. In view of these difficulties, we turned again to HEK293TN cells. As reported in Pietraszewska-Bogiel et al. (2019), we measured robust EKARcyt signals in cells transiently expressing H1R and EKARcyt sensor but not in cells expressing only the reporter (Fig. 5D). H1R-mediated EKARcyt signal was efficiently abolished upon Gα_q_ inhibition, although we measured weak and transient EKARcyt signals in a subset of FR900359-inhibited cells. Overexpression of either ßarr1-mCh or ßarr2-mCh diminished EKARcyt signal by more than 50%. On the contrary, EKARcyt signals in cells co-expressing H1R-p2A-mCh and EKARcyt that were first transfected with ctrl siRNA, ß-arr1 siRNA, ß-arr2 siRNA or simultaneously ß-arr1 and ß-arr2 siRNA constructs were not statistically different (Fig. 5E).

As a reference, we characterized the effect of overexpression or ß-arrestin knockdown on AT1_A_R-mediated ERK1/2 activation in HEK293TN cells, as it was thoroughly characterized by several groups using conventional methods for detection of ERK1/2 phosphorylation (Luttrell et al. 2001, Tohgo et al. 2002, Wei et al. 2003, Ahn et al. 2004ab). By doing so, we tested the performance of both EKARcyt sensor (see also Pietraszewska-Bogiel et al. 2019) and siRNA constructs employed in our studies. We measured a robust AngII-induced EKARcyt signal in cells co-expressing AT1_A_R-p2A-mCh together with EKARcyt sensor but not in cells expressing only the reporter (Fig. 6A). Overexpression of either ß-arrestin isoform reduced the signal by more than 50%, whereas Gα_q_ inhibition with FR900359 resulted in significantly reduced and more transient activation of ERK1/2 (Fig. 6A). In agreement with previous reports, silencing of ß-arr2 (Wei et al. 2003, Ahn et al. 2004ab, Kim et al. 2005) expression reduced the signal, especially at the later phase (Fig. 6B). Conversely, simultaneous knockdown of both ß-arrestin isoforms resulted in 20-30% increase of the EKARcyt signal.

**Figure 6.**
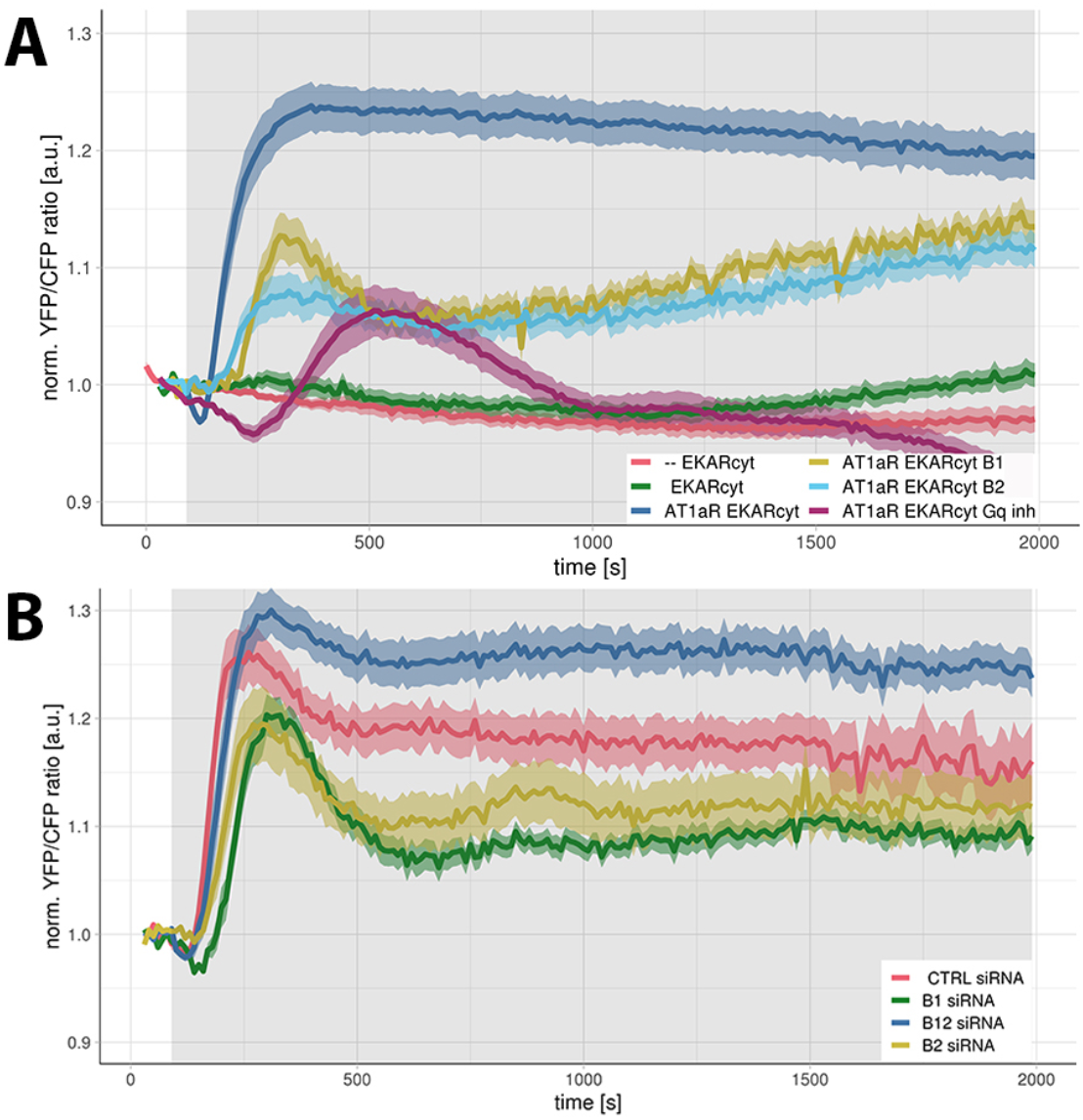
AT1_A_R-mediated ERK1/2 activation in HEK293TN cells. Time traces show the average ratio change of YFP/CFP fluorescence (±95% confidence intervals). Vehicle (--) or 1 μM AngII was added at 90 s; the grey boxes mark the duration of AngII stimulation. A) HEK293TN cells expressing EKARcyt sensor alone; co-expressing EKARcyt together with AT1_A_R-p2A-mCh and untreated or treated with Gα_q_ inhibitor (FR900359); co-expressing EKARcyt, AT1_A_R and ßarr1-mCh or ßarr2-mCh. B) HEK293TN cells co-expressing EKARcyt sensor together with AT1_A_R-p2A-mCh, previously transfected with ctrl siRNA, ß-arr1 siRNA, ß-arr2 siRNA or simultaneously ß-arr1 and ß-arr2 siRNA constructs.

Together, these results show that H1R activates ERK1/2 predominantly through heterotrimeric G proteins, whereas ß-arrestins act as negative regulators, desensitizing H1R-mediated ERK1/2 activation.

## DISCUSSION

Histamine is a major chemical mediator of allergic reactions whose action is mainly mediated by the H1R. However, up to date very little is known about the regulation of H1R-mediated signaling, especially by ß-arrestins. Here we employed pharmacological inhibition, overexpression and knockdown strategies, as well as dynamic measurements of signaling events in living cells to dissect the relative contributions of G proteins and ß-arrestins to H1R signaling.

We started by analyzing the character of H1R-ßarrestin interaction. As reported by Oakley and colleagues (2000), class A receptors bind β-arr2 with higher affinity than β-arr1, form transient complexes with β-arrestin, and undergo efficient recycling upon endocytosis. For example, ß2AR-βarrestin complex dissociates at or near the plasma membrane shortly after the formation of clathrin coated pits, excluding β-arrestin from receptor-containing endocytic vesicles (Oakley et al. 1999, Zhang et al. 1999). On the contrary, class B receptors bind both β-arrestin isoforms with similar high affinities and form stable receptor-arrestin complexes that persist as the receptor undergoes endosomal sorting, which slows recycling and promotes lysosomal degradation. Using similar quantification method as that employed by Oakley and colleagues (2000), we showed that H1R likely belongs to class A receptors. We also showed increased abundance of cytosolic ß-arrestin upon pyrilamine addition compared to the baseline levels which indicates that already prior histamine stimulation, a small portion of ß-arrestin protein exists in a bound state, presumably at the plasma membrane. This hypothesis is in agreement with the residual constitutive activity of H1R reported previously (Bakker et al. 2000).

Furthermore, using specific and efficient inhibitors for Gα_q_ (FR900359) and Gα_i_ (PTX), we were able to decouple both ß-arr1 and ß-arr2 recruitment upon H1R activation from either G_q_- or G_i_-mediated signaling. However, we have not performed dual inhibition experiments to conclude that the observed recruitment was achieved in “ zero G protein” background. At the same time, we noted diminished recruitment of both ß-arrestin isoforms in the absence of PKC activation (achieved with a specific PKC inhibitor, Ro31-8425), as well as more transient recruitment in the absence of G_q_ activation. Our results are largely consistent with the findings of Grundmann and colleagues (2018) who demonstrated ß-arr2 recruitment to D prostanoid receptor 2, orphan GPCR17 and (to a lesser extent) free fatty acid receptor 2 in the absence of active G proteins. Similarly, inhibition or lack of G_q/11_ was shown not to affect ß-arrestin-dependent internalization of free fatty acid receptor 4 and thyrotropin-releasing hormone (Yu & Hinkle 1999, Alvarez-Curto et al. 2016). The involvement of PKC in ß-arrestin recruitment upon H1R activation remains to be further elucidated but is in agreement with the role of ß-arrestins in desensitization of GPCRs phosphorylated by second messenger-activated kinases (Tóth et al. 2018, Luo et al. 2017), especially in the view of PKC-dependent phosphorylation of H1R (Fujimoto et al. 1999, Horio et al. 2004). Conversely, ß-arrestin recruitment to AT1_A_R and ß2AR was shown to be predominantly dependent on GRK-mediated phosphorylation, and not on PKC or cAMP-dependent protein kinases (Kim et al. 2008, Violin et al. 2008a). Similar to Violin and colleagues (2006), we were able to show the stimulatory effect of GRK2 and GRK6 on H1R-mediated recruitment of both ß-arrestins upon GRK overexpression in HeLa cells. Surprisingly, catalytically impaired mutant of GRK2, but not of GRK6, was still able to support the observed increase in ß-arrestin relocation. It has to be noted that the loss-of-function caused by L66A E514A double mutation was demonstrated for GRK5 (Baameur et al. 2010) but not for highly homologous GRK6. However, due to evolutionary conservation of catalytically important residues (including L66 and E514 in GRK5) found in all GRKs (Baameur et al. 2010), observations in GRK5 should carry over to other members of the GRK family. Supporting this notion, double mutations other than L66A E514A were shown to cause similar catalytic impairment of both GRK5 and GRK6 (Baameur et al. 2010). Regarding GRK2, the D110A mutation was reported to disrupt the Regulator of G protein Signaling (RGS) homology domain of GRK2, rendering GRK2 mutant incapable of relocation to activated Gα_q_ (Sterne-Marr et al. 2003). However, overexpression of GRK2[D110A K220R] mutant in HEK293 cells resulted in only 40-50% inhibition of histamine-induced inositol phosphate production (Iwata et al. 2005), which indicates its residual ability to inhibit PLC activation. Additionally, the lack of kinase activity of GRK2[K220R] mutant was not demonstrated (Kong et al. 1994). It would be interesting to find out if this partial activity is similarly observed with regard to the ability of GRK2[D110A K220R] mutant to phosphorylate GPCRs. Such hypothesized residual kinase function of this mutant could explain its ability to enhance ß-arrestin relocation similarly to WT GRK2.

Taken together, our results indicate the involvement of GRK2 and GRK6 in the H1R-mediated recruitment of ß-arrestins, at least upon their overexpression in HeLa cells. Our results agree with the finding that phosphorylation and desensitization of H1R in uterine myometrial cells and HEK293 cells is predominantly GRK2-mediated (Iwata et al. 2005). However, insight in the dependence of this recruitment on phosphorylation requires additional experiments.

ß-arrestins constitute important regulators of GPCR function, whose central role is to desensitize G protein-mediated signaling (Alvarez-Curto et al. 2016, Grundmann et al. 2018, Luttrell et al. 2018, reviewed in Gurevich & Gurevich 2019). At the same time, ß-arrestins can scaffold various downstream effectors and were shown to have a stimulatory effect on the selected aspects of GPCR signaling (Jean-Charles et al. 2017, Grundmann et al. 2018). To investigate this (putatively) complex ß-arrestin role in H1R-mediated signaling, we implemented both ß-arrestin overexpression and knockdown strategies to study their effect on three different signaling outcomes of H1R activation, i.e. PLCß, RhoA and ERK1/2 activation. Overexpression of either ß-arr1 or ß-arr2 in HeLa cells did not affect the immediate phase of histamine-induced activation of PLCß but promoted its desensitization at later time points. Conversely, knockdown of either ß-arrestin isoform in HEK293TN cells slowed down desensitization of histamine-induced PLCß activation. Previously, depletion of ß-arr2, but not ß-arr1, was shown to prolong H1R-stimulated Ca^2+^ response in uterine myometrial cells (Brighton et al. 2011). However, as regulation of GPCRs is likely to be highly dependent on the cell background, our results do not necessarily contradict those of Brighton and colleagues.

We also showed that both ß-arrestin isoforms function as desensitizers of G_q/11_-mediated activation of RhoA in HUVEC cells and ERK1/2 activation in HEK293TN cells. Previously, selective ß-arr2 depletion was shown to abolish, whereas depletion of ß-arr1 to augment H1R-stimulated ERK1/2 activation in uterine myometrial cells (Brighton et al. 2011). We found that overexpression of either ßarr-1 or ßarr-2 reduced both DORA-RhoA and EKARcyt signals by approximately 50%. Interestingly, we did not observe any effect of ß-arrestin knockdown on H1R-mediated activation of ERK1/2 in HEK293TN cells. On the contrary, knockdown of either ß-arr1 or ß-arr2 decreased, whereas simultaneous knockdown of both ß-arrestin isoforms increased AT1_A_R-mediated ERK1/2 activation in those cells. Because ß-arrestins both negatively regulate G protein-dependent ERK1/2 activation by promoting GPCR desensitization, and at the same time promote this activation by functioning as ligand-regulated scaffolds, deletion of both ß-arrestin isoforms can be expected to have a complex effect on signaling to ERK1/2. As stated by Grundmann and colleagues (2018), the lack of effect on the net ERK1/2 activation upon simultaneous ß-arr1 and ß-arr2 knockdown could reflect the balance between augmented G protein-dependent signaling, which would increase the ERK1/2 signal, and reduced scaffolding function of ß-arrestins, which would decrease it. Conversely, enhanced AngII-induced ERK1/2 activation in cells depleted for both ß-arrestin isoforms would indicate that the effects of ß-arrestins on desensitization of G protein-dependent ERK activation overrode their potential direct contributions to ERK activation. The observed differences between the effect of ß-arrestin knockdown on H1R- and AT1_A_R-mediated ERK1/2 activation could reflect different stability of receptor-ßarrestin complexes formed (i.e. transient vs stable).

Finally, we report complete abolishment of H1R-mediated ERK1/2 activation in HeLa, HUVEC and HEK293TN cells incubated with FR900359, indicating strong, if not complete, dependence of this response from G_q_ activation. Our data on H1R-mediated ERK1/2 activation in HeLa cells agrees with a previous report (Jain et al. 2016), with regard to kinetics, PTX insensitivity and Gq-dependence. However, despite using similar reporters for ERK1/2 activation as Jain and colleagues (2016), we did find histamine-induced activation of the nuclear ERK1/2 pool, as indicated by EKARnuc signals. At the moment, we cannot explain this discrepancy in results. Histamine-induced up-regulation of H1R expression could be one of the outcomes of this nuclear ERK1/2 activity (Mizuguchi et al. 2011). Total dependence of ERK1/2 activation on G protein has recently been reported for a panel of GPCRs belonging to different G protein-coupling groups (Alvarez-Curto et al. 2016, Grundmann et al. 2018, see also Gurevich & Gurevich 2018).

Taken together, our results provide new insights into the role of ß-arrestins in H1R-mediated signaling and are a good starting point for future studies on ERK signaling dynamics triggered by histamine.

## MATERIALS and METHODS

### Constructs

Plasmids encoding HsH1R-mCherry and HsH1R-p2A-mCherry are described in van Unen et al. (2016a). The p2A viral sequence in the latter construct ensures that the mCherry is separated from the receptor protein during translation, generating untagged receptor and free RFP that reports on receptor translation levels. Plasmid encoding RnAT1_A_R-mVenus was a kind gift from Peter Várnai (Semmelweis University, Hungary). The AT1_A_R sequence was cloned into pN1-mCherry and pN1-p2A-mCherry (Clontech) using HindIII and AgeI sites. Untagged H1R and AT1_A_R, used for co-expression experiments with EKARcyt sensor and ßarr-mCherry were generated by PCR amplification and restriction cloning using reverse primers with stop codons introduced immediately after the H1R or AT1_A_R sequence. Plasmids encoding Hsß2AR-mCFP (plasmid #38260), Rnßarr1-mYFP (plasmid #36916) and Rnßarr2-mYFP (plasmid #36917) were purchased from addgene.org. Using PCR amplification and restriction cloning, the ß2AR sequence was cloned into pN1-mCherry to generate ß2AR-mCherry construct. Using PCR amplification and restriction cloning, the ß-arrestin sequences were cloned into pC1 or pN1 (Clontech) to generate, respectively, ß-arrestin fused at their N-terminus or C-terminus with a fluorescent protein (FP): mTurquoise2, sYFP2 or mCherry. The ßarr1-FP and ßarr2-FP constructs were generated using KpnI and AgeI sites in pN1-FP, FP-ßarr1 and FP-ßarr2 constructs were generated using BspEI and HindIII sites in pC1-FP (stop codon was introduced in the reverse primers for ß-arrestin sequence). Plasmids encoding Lck-mVenus (Addgene plasmid #84337; van Unen et al., 2016b) and the DORA-RhoA sensor (Unen et al., 2015) were reported before. Cerulean-Venus versions of the EKARcyt (plasmid #18679) and EKARnuc (plasmid #18681) biosensors (Harvey et al. 2008) were purchased from addgene.org. Plasmid encoding BtGRK2-eCFP was a kind gift from Jeffrey L. Benovic (Thomas Jefferson University, Philadelphia) and HsGRK6 (plasmid #32693) construct was purchased from addgene.org. The GRK2 sequence was cloned into pN1-sYFP2 vector (Clontech) using NheI and AgeI sites. Plasmid encoding GRK6-sYFP2 was generated by cloning GRK6 sequence into pN1-sYFP2 vector using PCR amplification and EcoRI and AgeI sites. Mutations D110A K220R and L66A E514A were introduced into, respectively, GRK2 and GRK6 sequences using site-directed mutagenesis; after the mutations were confirmed with sequencing, GRK2 [D110A K220R] and GRK6 [L66A E514A] sequences were cloned into pN1-sYFP2. All constructs used in this study will be deposited at www.addgene.org.

### Reagents

Histamine, angiotensin II (AngII), pyrilamine (PY), pertussis toxin (PTX), and Ro31-8425 were purchased from Sigma Aldrich/Merck. The specific Gα_q_ inhibitor, FR900359 (Ubo-QIC), was purchased from the University of Bonn (http://www.pharmbio.uni-bonn.de/signaltransduktion). Cells in microscopy medium were incubated with 1 μM FR900359 at least 10 minutes before the start of the measurements. Such treatment was sufficient to efficiently abrogate histamine-induced G_q_ activation and PIP2 degradation in HeLa and HEK293TN cell transiently expressing H1R (data not shown). Ro31-8425 was added to cells (in microscopy medium) at least 25 minutes before the measurements at a concentration of 10 μM. Experiments with PTX inhibitor were carried out using cells cultured in serum free DMEM and incubated o/n with 100 ng/μL PTX.

### Cell culture & sample preparation

HeLa cervix carcinoma cells (American Tissue Culture Collection: Manassas, VA, USA) and human embryonic kidney (HEK) 293TN cells (System Biosciences, LV900A-1) were cultured using Dulbecco’ s Modified Eagle Medium (DMEM) supplied with glutamax, 10% FBS, penicillin (100 U/ml) and streptomycin (100 ug/ml), and incubated at 37°C and 5% CO_2_. All cell culture regents were obtained from Invitrogen (Bleiswijk, NL). Cells were transfected in a 35 mm dish holding a glass coverslip (24 mm ∅, Menzel-Gläser, Braunschweig, Germany), using polyethyleneimine (3 μL of PEI:1 μL of DNA) according to the manufacturer’ s protocol. For each transfection, we used 500 ng of receptor (H1R, AT1_A_R or ß2AR)-carrying plasmid. Other plasmids were transfected at: 100 ng (ß-arr1 fusion constructs), 150 ng (ß-arr2 fusion constructs), 200 ng (Lck-mVenus, EKARnuc), 250 ng (EKARcyt, mTQ2-PHM), 300 ng (DORA-RhoA reporter, mTQ2-Rab7 fusion).

Human umbilical vein endothelial cell (HUVEC) cell culture and transfection was performed as described in Reinhard et. al. (2016). Cells were transfected using electroporator according to the manufacturer’ s protocol. 1 μg cDNA of EKARcyt or DORA-RhoA reporter was used per transfection.

Samples were imaged one day after transfection: coverslips were mounted in an Attofluor cell chamber (Invitrogen, Breda, NL) and submerged in 1 mL microscopy medium (20 mM HEPES, pH=7.4, 137 mM NaCl, 5.4 mL KCl, 1.8 mM CaCl_2_, 0.8 mM MgCl_2_, and 20 mM glucose; Sigma Aldrich/Merck).

### Silencing of ß-arrestin expression

DharmaFECT1 transfection reagent, as well as ctrl, ß-arr1 and ß-arr2 siRNA constructs were purchased from Dharmacon: Non-targeting pool 5 nmol D-001810-10-05 (ctrl siRNA), and SMARTpool 5nmol L-011971-00-0005 (ß-arr1 siRNA) and L-007292-00-0005 (ß-arr2 siRNA) constructs. Thirty % confluent HEK293TN cells were transfected with 2 μL of siRNA constructs (prepared at 2 μM concentration) using DharmaFECT1 and Opti-MEM according to the manufacturer’ s protocol.

### Immunoblotting

Cells were lysed in 20 μl Passive Lysis Buffer (Promega) and protein concentration was measured using the Pierce BCA Protein Assay Kit (Thermo Fisher Scientific). Equal amounts of protein (4 μg) were run on a 12% SDS-PAGE gel. Proteins were transferred to a nitrocellulose membrane (Bio-Rad) and blocked with Odyssey Blocking Buffer (LI-COR Biosciences, diluted 1:1 in TBS prior to use). Primary mouse antibody directed against ß-arrestin (Cat#610550 BD Transduction Lab, 1:300 dilution; recognizes both ß-arrestin isoforms) and rabbit antibody directed against Hsp90 (Cat#610418 BD Transduction Lab, 1:1000 dilution) were diluted in blocking buffer supplemented with 0.1% Tween-20 (TBS-T). Staining was performed overnight at 4°C. Membranes were washed in TBS-T followed by 2 hour-incubation with secondary antibodies: IRDye 680LT (Cat#926-68021) or IRDye 800CW (Cat#926-32212) (LI-COR), diluted at 1:20000 in TBS-T). Membranes were washed in TBS-T and incubated in TBS prior to scanning at 700 nm and 800 nm using an Odyssey Fc Imaging System (LI-COR Biosciences). Image Studio™ Lite 4.0 software (LI-COR Biosciences) was used to visualize the data.

### Confocal and wide-field microscopy

Imaging of subcellular localization and relocation experiments on the confocal microscope were performed as described in van Unen et al. (2016b). Ratiometric FRET measurements were performed on a previously described wide-field microscope (van Unen et al. 2015). Typical exposure time was 100 ms, and camera binning was set to 4×4. Fluorophores were excited with 420/30 nm light and reflected onto the sample by a 455DCLP dichroic mirror. CFP emission was detected with a BP470/30 filter and YFP emission was detected with a BP535/30 filter by rotating the filter wheel. RFP was excited with 570/10 nm light reflected onto the sample by a 585 dichroic mirror, and RFP emission was detected with a BP620/60 nm emission filter. All acquisitions were corrected for background signal and bleedthrough of CFP emission in the YFP channel. All experiments were performed at 37°C.

For ratiometric relocation measurements, H1R-p2A-mCherry construct was used (in place of H1R-mCherry) in order to correct for changes in cell height by measuring the relative CFP/RFP fluorescence ratio). Ratiometric relocation measurements were performed by using the above CFP excitation and emission settings, and the RFP was imaged as follows: RFP was excited with 570/10 nm light reflected onto the sample by a 585 dichroic mirror, and RFP emission was detected with a BP620/60 nm emission filter. To confirm the expression of GRK-sYFP2 fusions, YFP fluorescence was measured as follows: YFP was excited with 500/30 nm light reflected onto the sample by a 515DCXR dichroic mirror, and YFP emission was detected with a BP535/30 nm emission filter. All acquisitions were corrected for background signal and bleedthrough of CFP emission in the YFP channel. Bleedthrough of CFP in the RFP channel was negligible.

### Image analysis and data visualization

ImageJ (National Institute of Health) was used to analyze the raw microscopy images. Background subtractions, bleedthrough correction and calculation of the normalized ratio per time point per cell were done in Excel (Microsoft Office). All plots were prepared with the PlotTwist web app (Goedhart 2019). Plots show the average response as a thicker line and a ribbon for the 95% confidence interval around the mean.

## Author contributions

APB designed and performed experiments, analyzed the data, and wrote the manuscript. JG participated in study design, data interpretation, and data visualization. All authors approved the final manuscript.

## Acknowledgments & Funding

We thank Peter Várnai (Semmelweis University, Hungary) and Jeffrey L. Benovic (Thomas Jefferson University, Philadelphia, the USA) for providing the constructs, and Nathalie Reinhard (University of Amsterdam, the Netherlands) for providing assistance with HUVEC maintenance and transfection, and Nika Heijmans and Katrine Wiese (University of Amsterdam, the Netherlands) for providing assistance with immunoblotting. A Pietraszewska-Bogiel was supported by NanoNextNL, a micro and nanotechnology consortium of the Government of the Netherlands and 130 partners.

## SUPPLEMENTARY DATA

**Supp. Figure 1.**
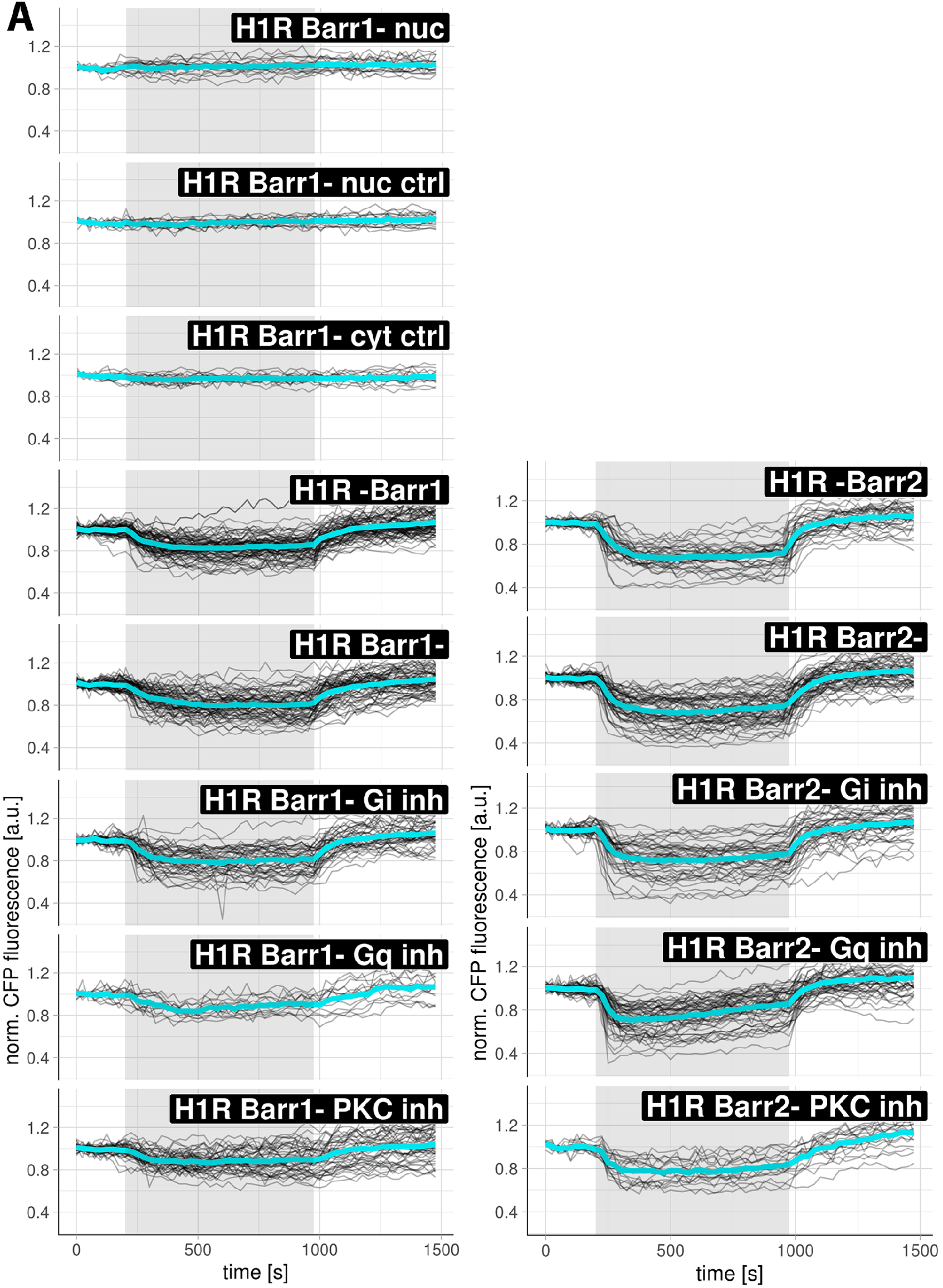
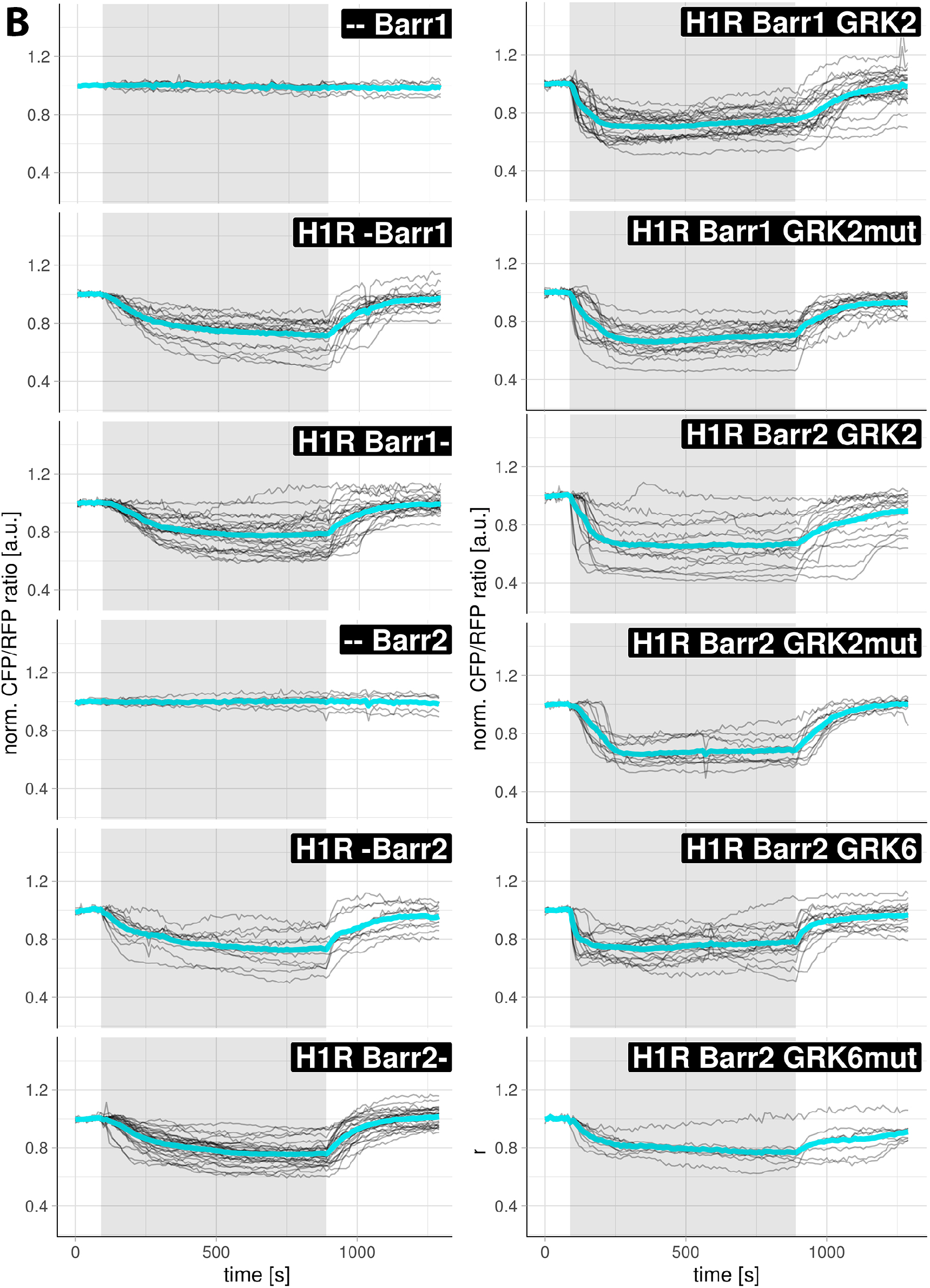
Quantification of ß-arrestin recruitment in HEK293TN cells. A) Confocal measurements of ß-arrestin relocation. Black lines show the CFP fluorescence in individual cells (measured in cytosol, except for the two top most graphs that show measurements of ß-arr1 abundance in the nucleus), and the average fluorescence is shown as a thicker cyan line. Black boxes specify the constructs (co-)expressed in cells. Left: HEK293TN cells co-expressing: H1R-mCh and mTQ2-ßarr1 or ßarr1-mTQ2 and untreated, treated with Gα_i_ inhibitor (PTX), Gα_q_ inhibitor (FR900359) or PKC inhibitor (Ro31-8425). Right: HEK293TN cells co-expressing: H1R-mCh and mTQ2-ßarr2 or ßarr2-mTQ2 and untreated, treated with PTX, FR900359 or Ro31-8425. 100 μM histamine was added at 200 s, 10 μM PY was added at 950 s. Grey boxes mark the duration of histamine stimulation. B) Wide-field measurements of ß-arrestin relocation. Black lines show the ratio change of CFP/RFP fluorescence in individual cells (measured in cytosol, except for the two top most graphs that show measurements of ß-arr1 abundance in the nucleus), and the average ratio change is shown as a thicker cyan line. Left: HeLa cells expressing: ßarr1-mTQ2 alone; co-expressing H1R-p2A-mCh together with ßarr1-mTQ2, mTQ2-ßarr1; ßarr2-mTQ2 alone; co-expressing H1R-p2A-mCh together with ßarr2-mTQ2 or mTQ2-ßarr2 and stimulated with vehicle (--) or histamine (100 μM histamine was added at 90 s and 10 μM PY at 890 s). Unless otherwise stated, the grey boxes mark the duration of histamine stimulation. Right: HeLa cells co-expressing: H1R-p2A-mCh, ßarr1-mTQ2 and GRK2-sYFP2; H1R-p2A-mCh, ßarr1-mTQ2 and GRK2[D110A K220R]-sYFP2; H1R-p2A-mCh, ßarr2-mTQ2 and GRK2-sYFP2; H1R-p2A-mCh, ßarr2-mTQ2 and GRK2[D110A K220R]-sYFP2; H1R-p2A-mCh, ßarr2-mTQ2 and GRK6-sYFP2; H1R-p2A-mCh, ßarr2-mTQ2 and GRK6[L66A E514A]-sYFP2. 100 μM histamine was added at 90 s and 10 μM PY at 890 s; the grey boxes mark the duration of histamine stimulation.

**Supp. Figure 2.**
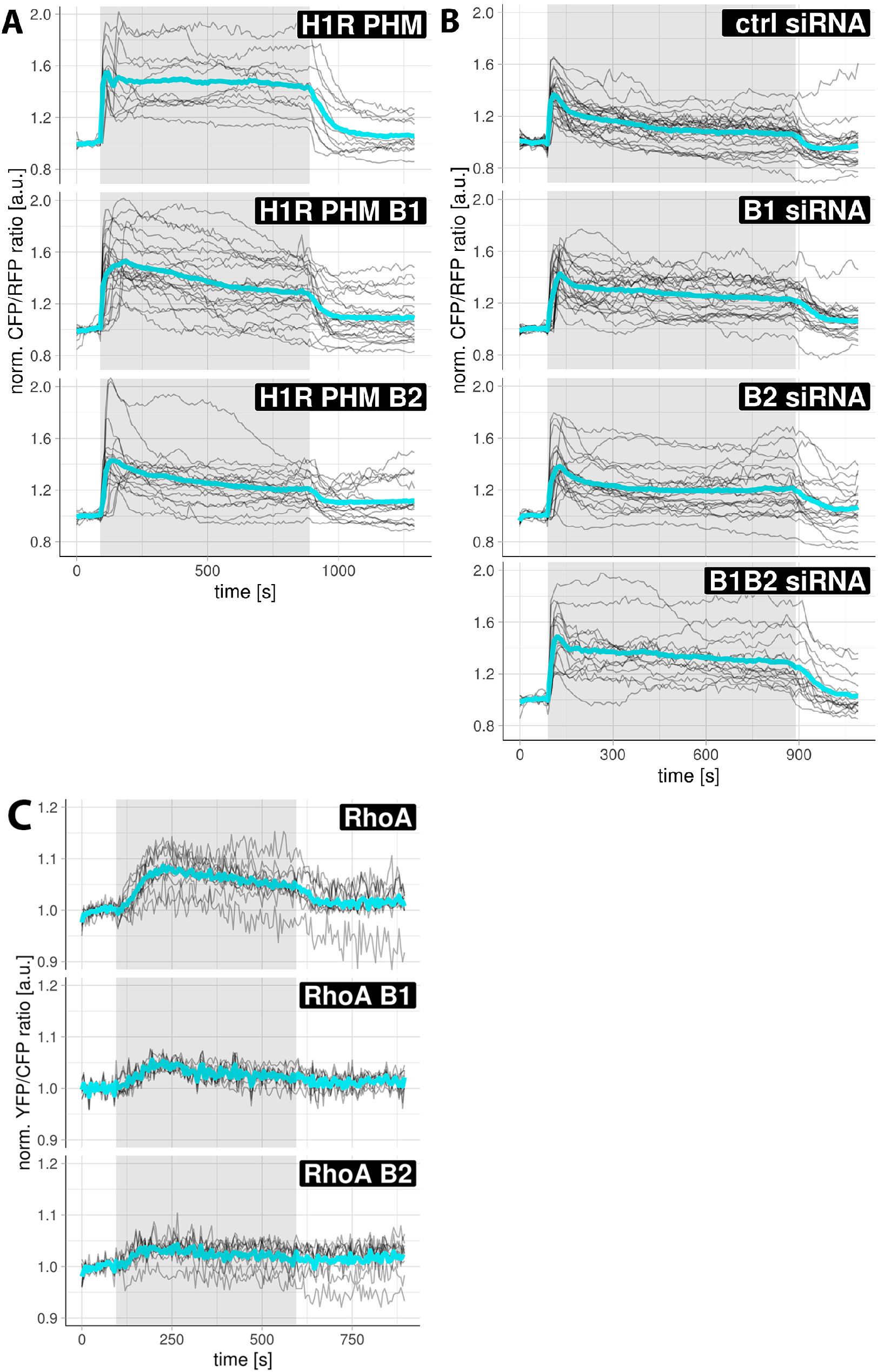
Role of ß-arrestins in G_q_-mediated signaling downstream of H1R. Black lines show the ratio change of YFP/CFP fluorescence in individual cells, and the average ratio change is shown as a thicker cyan line. Black boxes specify the constructs (co-)expressed in cells. A-B) The effect of ß-arrestin overexpression and knockdown on histamine-induced PLC activity. 100 μM histamine was added at 90 s and 10 μM PY at 890 s; grey boxes mark the duration of histamine stimulation. A) HeLa cells co-expressing: H1R-p2A-mCh together with PIP2 reporter (mTQ2-PHM); H1R-p2A-mCh, mTQ2-PHM and ßarr1-sYFP2 or ßarr2-sYFP2. B) HEK293TN cells expressing H1R-p2A-mCh and mTQ2-PHM, that were first transfected with ctrl siRNA, ßarr1 siRNA, ßarr2 siRNA or simultaneously ßarr1 and ßarr2 siRNA constructs. C) The effect of ß-arrestin overexpression on histamine-induced RhoA activation. HUVEC cells expressing DORA-RhoA reporter (RhoA) alone or together with either ßarr1-mCherry or ßarr2-mCherry. 100 μM histamine was added at 95 s and 10 μM PY at 595 s; grey boxes mark the duration of histamine stimulation.

**Supp. Figure 3.**
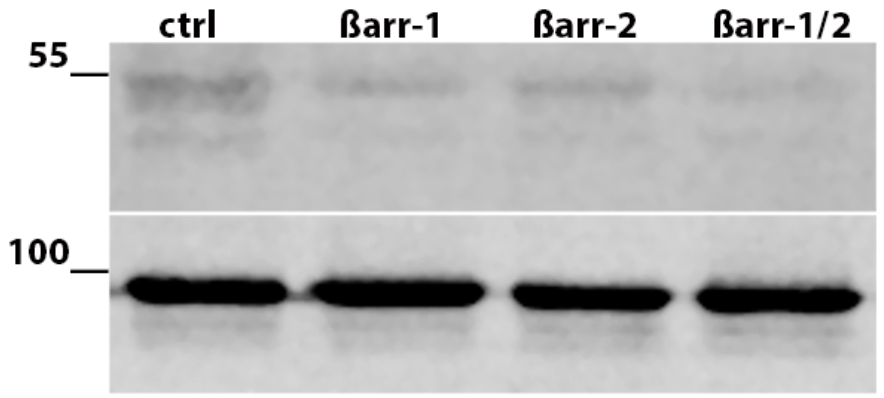
Efficiency of silencing of ß-arr1 and/or ß-arr2 expression in HEK293TN cells. HEK293TN cells transfected with: ctrl siRNA, ß-arr1 siRNA, ß-arr2 siRNA or simultaneously ß-arr1 and ß-arr2 siRNA constructs. Two days later, whole protein was isolated and separated on an SDS gel. ß-arrestin (upper panel) was detected using purified anti-ß-arrestin Ab that recognizes both isoforms (upper band: ß-arr1, lower band: ß-arr2). Equal protein loading was confirmed using purified anti-Hsp90 Ab (lower panel).

**Supp. Figure 4.**
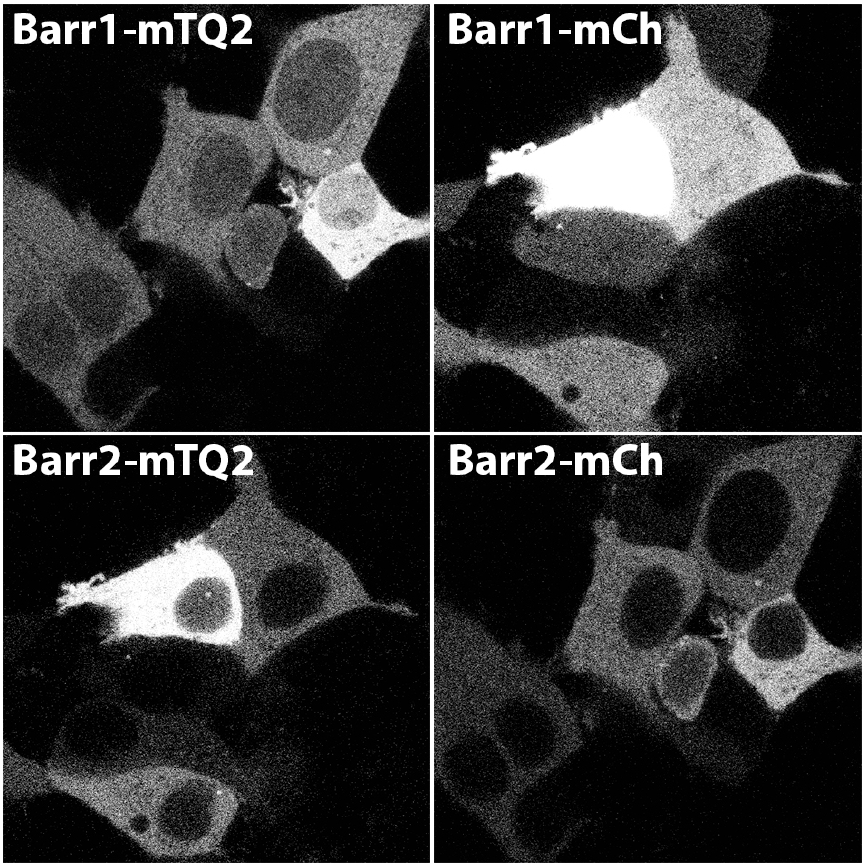
Subcellular localization of ß-arr1 and ß-arr2 fusion proteins in HEK293TN cells. Confocal images depicting subcellular localization of ßarr1-mTQ2, ßarr1-mCh, ßarr2-mTQ2 and ßarr2-mCh in HEK293TN cells. The size of the images is 70 × 70 μm.

**Supp. Figure 5.**
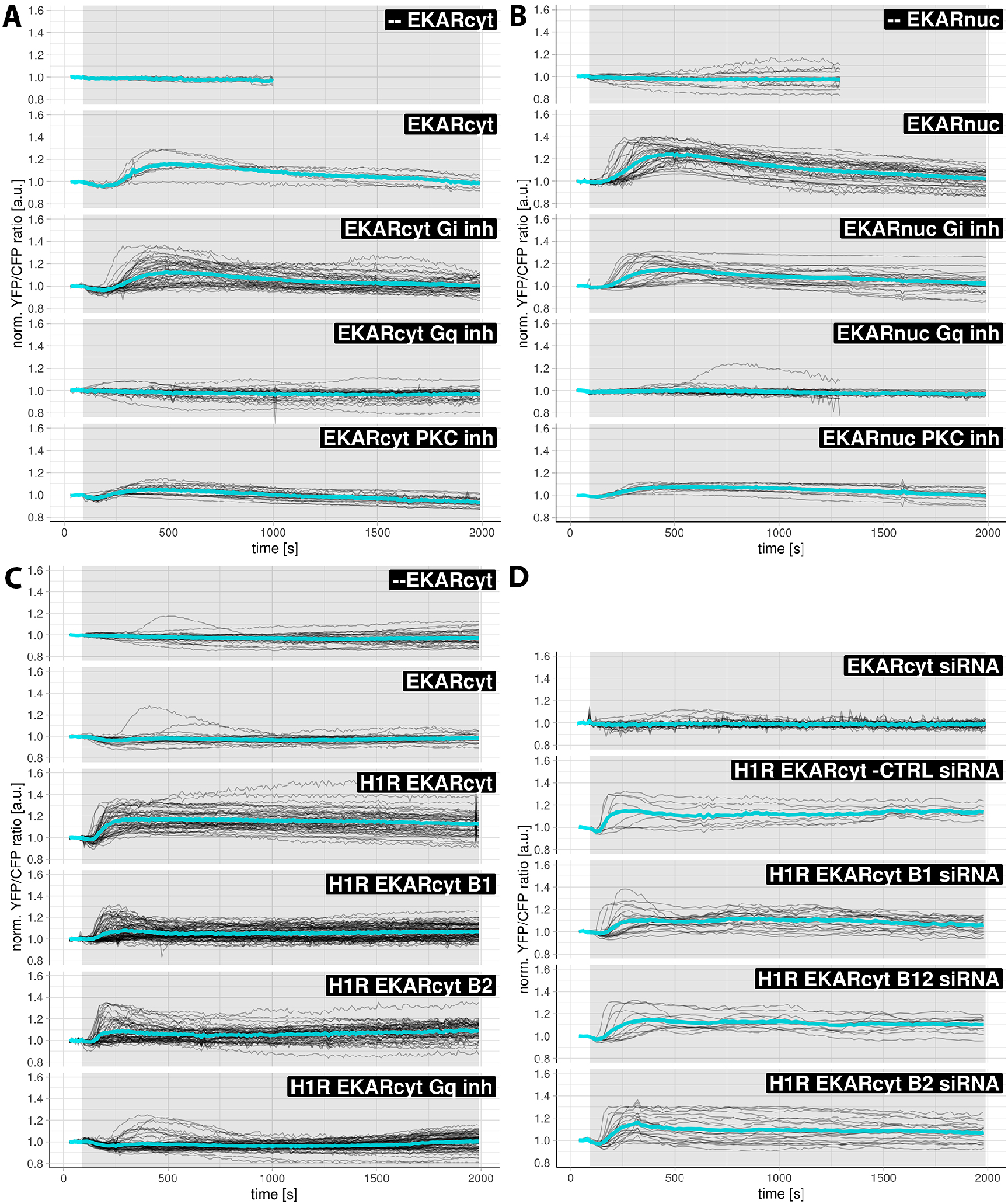
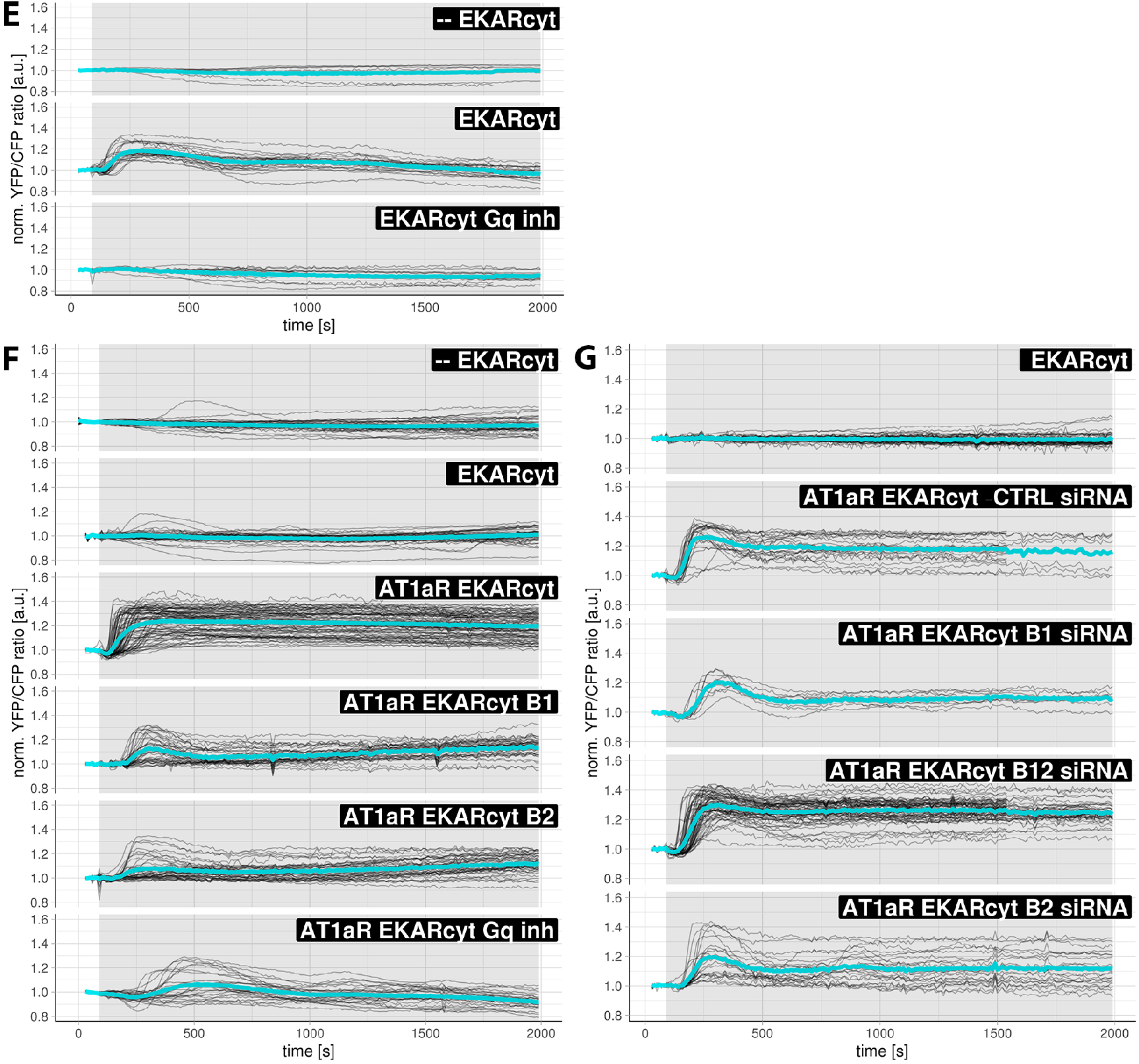
Histamine- and AngII-induced ERK1/2 activation. Black lines show the ratio change of YFP/CFP fluorescence in individual cells, and the average ratio change is shown as a thicker cyan line. Vehicle (--) or 100 μM histamine was added at 90 s; the grey boxes mark the duration of histamine stimulation. A-B) HeLa cells expressing EKARcyt (A) or EKARnuc (B) sensor and untreated, treated with Gα_i_ inhibitor (PTX), Gα_q_ inhibitor (FR900359) or PKC inhibitor (Ro31-8425), C-D) HEK293TN cells: C) expressing EKARcyt sensor alone (N=26); co-expressing H1R-p2A-mCh and EKARcyt and untreated or treated with Gα_q_ inhibitor (FR900359); co-expressing H1R, EKARcyt and ßarr1-mCherry or ßarr2-mCherry; D) co-expressing H1R-p2A-mCh and EKARcyt that were first transfected with ctrl siRNA, ß-arr1 siRNA, ß-arr2 siRNA or simultaneously ß-arr1 and ß-arr2 siRNA constructs. E) HUVEC cells expressing EKARcyt and untreated or treated with FR900359, F-G) HEK293TN cells: F) expressing EKARcyt sensor alone; co-expressing AT1_A_R-p2A-mCh and EKARcyt and untreated or treated with FR900359; co-expressing AT1_A_R-p2A-mCh, EKARcyt and ßarr1-mCh or ßarr2-mCh; G) co-expressing AT1_A_R-p2A-mCherry and EKARcyt that were first transfected with ctrl siRNA, ß-arr1 siRNA, ß-arr2 siRNA or simultaneously ß-arr1 and ß-arr2 siRNA constructs.

**Supp. Movies. ß-arrestin recruitment to plasma membrane upon H1R activation in HEK293TN cells**.

1-4) Wide-field recordings of subcellular localization of ßarr1-mTQ2 (1), mTQ2-ßarr1 (2), mTQ2-ßarr2 (3) or ßarr2-mTQ2 (4) co-expressed together with H1R-p2A-mCh in HEK293TN cells. 100 μM histamine was added at 90 s and 10 μM PY at 890 s. Time interval is 25 s. The size of the images is 105 × 141 μm.

5-8) Confocal recordings of subcellular localization of ßarr1-mTQ2 (5), mTQ2-ßarr1 (6), mTQ2-ßarr2 (7) or ßarr2-mTQ2 (8) co-expressed together with H1R-mCh in HEK293TN cells. 100 μM histamine was added at 225 s and 10 μM PY at 975 s. Time interval is 25 s. The size of the images is 60 × 60 μm.

9-10) Confocal recordings of subcellular localization of ßarr1-mTQ2 (9) or ßarr2-mTQ2 (10) coexpressed together with ß2AR-mCh in HEK293TN cells. 10 μM isoproterenol was added at 225 s and 50 μM propranolol at 975 s. Time interval is 25 s. The size of the images is 60 × 60 μm.

11-12) Confocal recordings of subcellular localization of ßarr1-mTQ2 (11) or ßarr2-mTQ2 (12) coexpressed together with AT1_A_R-mCh in HEK293TN cells. 1 μM AngII was added at 225 s. Time interval is 25 s. The size of the images is 60 × 60 μm.

